# A Mechanical Logic for Bacterial Navigation

**DOI:** 10.1101/2025.05.27.656390

**Authors:** Rui Zeng, Stefano Sacanna, Saumya Saurabh

## Abstract

Bacterial motility in unconfined liquids is well understood, but many species inhabit crowded, three-dimensional environments where physical constraints shape navigation. Here, we show that the aquatic bacterium *Caulobacter crescentus* employs backward swimming to overcome confinement, using a force-sensitive mechanism to control directional switching. Using colloidal “µ-Traps” to confine single cells within spherical volumes accessible through a narrow aperture, we reveal that cells engage backward motility to enter pores, explore boundaries, and escape confinement. This directional switching is gated by mechanical load: moderate forces on the flagellum promote backward motion, while excessive load suppresses it, preventing futile effort. This tunable, load-sensitive control offers direct evidence for a mechanical logic that governs motility: backward movement is activated when advantageous and disengaged when not. These findings highlight how physical forces influence microbial navigation and adaptation in complex environments.

## Introduction

Bacterial motility is fundamental to survival, influencing processes ranging from colonization and nutrient acquisition to pathogenesis (*1–3*). Classic studies of *Escherichia coli* motility in liquid and two-dimensional (2D) environments have established the foundational “run-and-tumble” model, in which directional changes are controlled by the coordinated action of multiple flagella (*1, 4*). While this paradigm has been instrumental in shaping our understanding of bacterial navigation (*5*), it primarily reflects the behavior of peritrichously flagellated species in relatively unstructured environments. Many other bacteria, including those with a single polar flagellum, must contend with physically complex habitats such as soil, host tissues, and biofilms, settings where confinement, surface contact, and spatial heterogeneity may require distinct motility strategies (*6–8*). These diverse habitats and architectures raise a longstanding question: why and how do bacteria swim backward?

To navigate these crowded, three-dimensional (3D) environments, many bacteria including *monotrichous* species, employ alternative motility modes tailored to physical constraints. The aquatic bacterium *Caulobacter crescentus* (*9*), for instance, integrates reversible flagellar rotation (*10*) with pili-mediated surface sensing (*11*) to dynamically adapt its movement (*9, 12–14*). Unlike *E. coli*, which is biased toward forward propulsion, *Caulobacter* frequently reverses direction, which is controlled by the chemotaxis response regulator CheY and a family of Cle proteins that modulate the flagellar switch complex (*15*).

Cryo-electron tomography has resolved structural rearrangements associated with forward and reverse states of the motor (*16*), and biophysical studies have explored whether asymmetric torque generation underlies the switch. While early work reported similar forces in both directions (*17*), later studies revealed higher torque in the reverse mode (*18*), suggesting functional asymmetry may confer a mechanical advantage in specific contexts. Because these findings emerged from systems with differing geometries, direct comparisons remain difficult. This variability has complicated efforts to resolve the underlying physical cues that regulate directional switching.

Although force-dependent motor switching has been proposed as a mechanism to reorient under load (*19–21*), its role in mediating bacterial behavior under realistic physical constraints remains poorly understood. The lack of suitable 3D experimental platforms has been a major obstacle: while 2D substrates dominate motility studies, most 3D imaging is performed in open, unconfined volumes that fail to capture key physical features of natural environments—such as tight curvature, narrow apertures, and microscale pores (*22, 23*). Even cutting-edge systems such as optical traps and nanofabricated channels offer limited insight into how cells transition between motility states within spatially complex microenvironments (*24, 25*).

Here, we investigate how backward motility emerges as a context-dependent strategy for navigating confined environments. Using colloidal “µ-traps”, microscale spherical compartments with a single aperture, and 3D single cell tracking, we discover a force-sensitive mechanism that regulates directional switching. The prevalence of such switching varies across physical contexts, suggesting an intrinsic ability to tune motility in response to spatial and mechanical cues. This dynamic modulation enables *monotrichous* bacteria convert physical signals into actionable motility responses, enabling rapid adaptation to microgeometric challenges in their surroundings. Understanding how physical features such as confinement and surface curvature influence bacterial behavior is particularly urgent for environmental and biomedical applications, where the ability to predict or control microbial distributions hinges on clarifying how cells respond to mechanical constraints in natural and engineered settings. (*8, 26, 27*).

## Results

### Functional role of backward swimming in navigating confinement

It has been speculated that the increased torque during backward motion could be advantageous for the escape of daughter cells after division (*18*). To assess whether reversible flagellar rotation confers a competitive advantage under physical confinement, we placed a filter paper with 1 µm pore-size on a nutrient-rich agarose plate for three minutes and subsequently added motile *Caulobacter* cells to the top surface (Fig. 1a). The filter paper formed a fibrous, porous mesh reminiscent of soil and aquatic microenvironments, where bacteria often navigate dense 3D structures. Scanning electron microscopy (SEM) confirmed a highly entangled architecture with micron-scale pores (Fig. S1), presenting a physiologically relevant barrier that cells must traverse to access nutrients. Wild-Type (WT) *Caulobacter crescentus*, which can reverse their single flagellum, readily traversed the filter and formed visible colonies underneath (Fig. 1b). In contrast, a mutant devoid of CheR chemotaxis proteins and unable to reverse (*28*) —referred to as Forward-Only (FO) in this study—exhibited markedly reduced passage and minimal growth on the far side of the filter (Fig. 1b). In the absence of the filter paper, the two strains showed similar fitness and colonization behavior (Fig. S2), suggesting no role of the backward motion in faster growth, division, or proliferation (*18*). These observations demonstrate that backward motion is critical for navigating micro-porous barriers and thereby securing access to nutrient-rich regions. However, these findings also raise two key mechanistic questions: How does backward motion confer an advantage in confined environments? And what triggers the switch to backward motility in response to physical constraints? Addressing these questions is challenging because extant methods for studying bacterial motility rely on open, 2D substrates or microfluidic channels, which fail to reproduce the geometric complexity and confinement present in natural environments. Real-time observation of bacterial behavior in physically constrained 3D spaces requires new, minimally perturbative approaches that preserve long-term cell viability and allow for precise measurements of motility dynamics.

**Figure 1:**
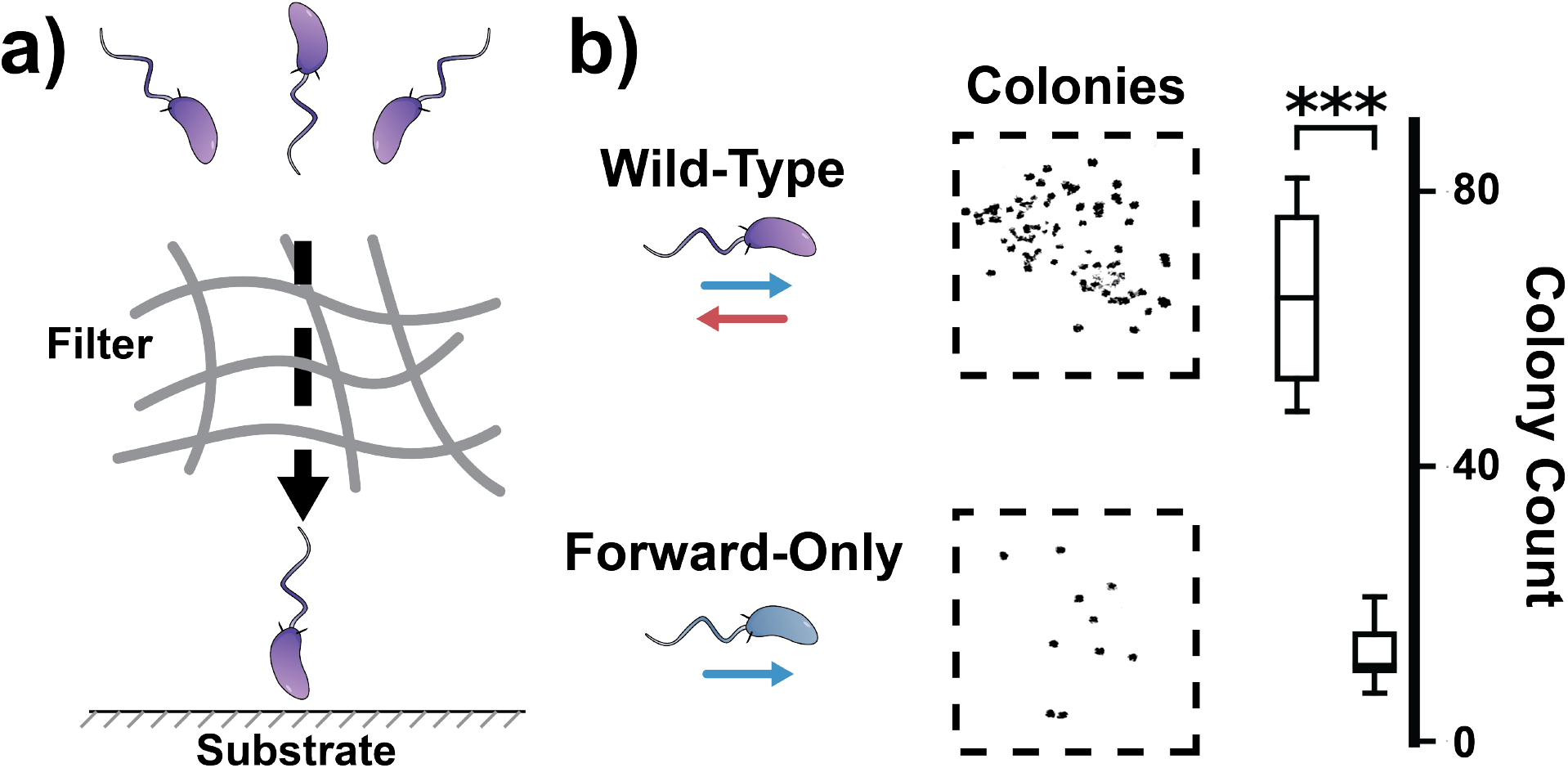
Backward motility enables access to nutrients through a porous mesh barrier. **a)** Schematic of the filter assay. A 1 µm pore-size filter paper is briefly placed on a nutrient-rich agarose plate, and motile *Caulobacter* cells are added on top. The fibrous mesh creates a 3D physical barrier between the cells and the underlying nutrients. Only bacteria that successfully traverse the micron-scale pores can access the nutrient-rich substrate and form colonies. **b)** Wild-Type *Caulobacter*, capable of reversible flagellar rotation, readily penetrates the mesh and colonizes the region beneath the filter. In contrast, a mutant lacking backward motility shows significantly reduced colony formation. Right: Quantification of colony counts beneath the filter from six independent experiments (box plot; *** *p <* 0.001). These results demonstrate that backward motility is essential for navigating structured 3D barriers and securing access to nutrient-rich microenvironments.

### Colloidal µ-Traps reveal single-cell dynamics under spatial confinement

While the filter assay reveals that backward motility enhances access to nutrients in structured environments, it leaves unanswered how individual cells physically maneuver, reorient, or adapt their behavior within confined spaces. We therefore applied colloidal µ-Traps (*29*), spherical shells (5–10 µm in diameter) with a single aperture (∼1 µm in diameter) designed to trap a single *Caulobacter* swimmer cell (Fig. 2a-b, Fig. S3). Brightfield (BF) and SEM images of these µ-Traps (Fig. S4) confirm their uniform geometry and narrow apertures. Once inside, cells (∼1.8 µm long, Fig. S4d) swim freely within the 3D volume for tens to hundreds of seconds. We combined highspeed BF microscopy with a custom 3D tracking algorithm. This approach exploits the axial phase shift in BF imaging, where cells above or below the focal plane exhibit distinct phase patterns to localize bacteria in all three dimensions and reconstruct their trajectories over time (Fig. 2c-d, Fig. S5-6, movie S1).

**Figure 2:**
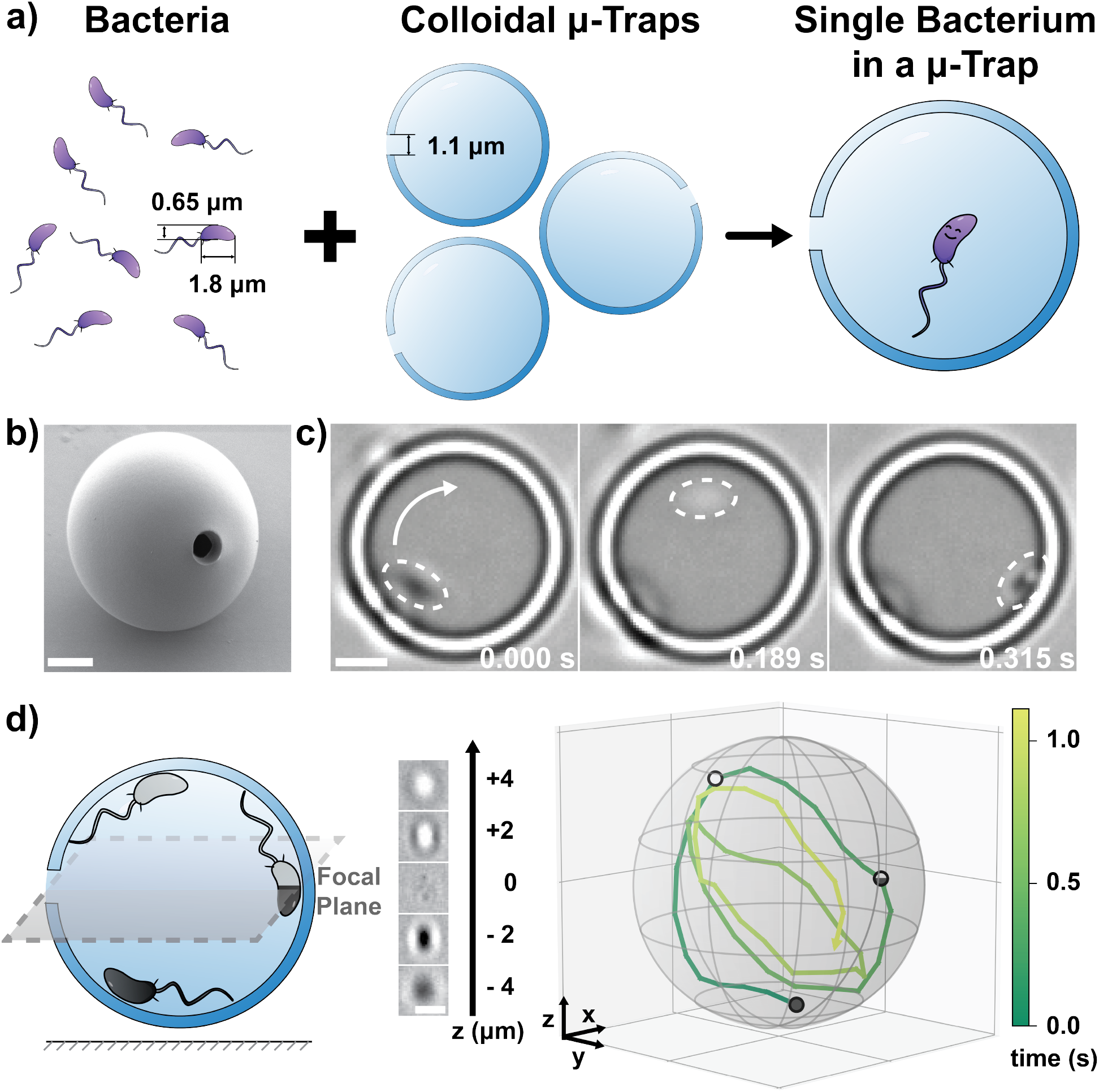
Colloidal µ-Traps enable 3D-resolved tracking of confined bacterial motion. **a)** Schematic of the experimental workflow: motile *Caulobacter* cells are introduced into a suspension containing colloidal µ-Traps. **b)** Scanning electron micrograph of a µ-Trap, a spherical hollow compartment with a single ∼1.1 µm aperture. **c)** Time-lapse brightfield images showing a trapped *Caulobacter* cell executing a circular trajectory near the compartment wall (dashed line indicates cell body). The cell remains motile within the confined volume. **d)** Schematic of axial localization by brightfield tracking. Cell z-position is inferred by quantifying phase-dependent diffraction pattern shape (insets, middle), enabling high-speed 3D tracking within a single trap. Grid size: 4 µm. Scale bars: 2 µm.

Within the confined geometry of the µ-Trap, cells displayed a repertoire of distinct 3D motion modes (Fig. S7, movie S2). These include: (i) forward swimming aligned with the trap wall (Forward I) (Fig. S7a), (ii) forward swimming at an angle to the wall, resulting in localized circling (Forward II) (Fig. S7b), and (iii) backward swimming following a directional switch, marked by tight circling near the trap center (Backward mode) (Fig. S7c). Corresponding trajectories reveal that cells frequently switch between these modes, forming complex spatial paths that wrap around the µ-Trap interior. Quantitative analysis (Fig. S8) showed that the three motility modes differ in key parameters including swimming speed, distance from surface, track radius, and radial offset. For example, Forward I tracks follow smoother arcs close to the surface, whereas Backward tracks remain centralized with tighter radii and lower velocities. These metrics establish that each mode offers unique geometrical or mechanical advantages depending on the cell’s orientation and the constraints imposed by the trap.

To further confirm that motion modes corresponded to flagellar-driven transitions, we fluorescently labeled the flagella and verified orientation-reversal dynamics under confinement (see methods, Fig. S9a, movie S2). These experiments also revealed a strong wavelength dependence in phototoxicity: cells exposed to 488 nm illumination exhibited rapid motility loss, whereas those imaged under 561 nm retained activity, guiding our choice of illumination laser for subsequent analyses (Fig. S9b-c, movie S3). To rule out potential artifacts from the imaging system, we confirmed that surfactant treatment (Pluronic^®^ F108) had no significant effect on the speed distribution of either Wild-Type or Forward-Only cells (Fig. S10). Thus, the motility patterns we observed arise from intrinsic cell behavior in confinement rather than surface adhesion or optical perturbation. Together, these findings demonstrate that *Caulobacter* swimmer cells dynamically access context-specific strategies tailored to spatial geometry.

Unlike conventional two-dimensional assays that tether cells to flat substrates (*18,19*), the µ-Trap configuration maintains a confined yet unbounded 3D environment, enabling prolonged observation of free-swimming bacteria under controlled spatial constraints. The ability to reorient via multiple motion modes—each with distinct geometric profiles and speed characteristics—demonstrates a finely tuned motility program adapted to confinement. Cumulatively, µ-Traps provide a powerful platform for dissecting the physical logic underlying bacterial exploration in complex microenvironments.

### Backward swimming improves entry, exploration, and escape under confinement

Based on these observations, we next asked how *Caulobacter crescentus* physically navigates the tightest regions of the trap: specifically, how cells enter, explore, and exit through a single narrow aperture. Representative brightfield images (Fig. 3a,c) show a swimmer cell approaching the µ-Trap aperture and then leaving it after a period of confinement. Strikingly, backward motion was strongly favored during both events. After normalizing by the forward-backward swimming bias observed outside and inside the trap (∼34% backward swimmers outside u-Traps, Fig. 3b; ∼35% backward swimmers inside u-Traps, Fig. 4b), we found that backward entries and escapes occurred more frequently—69% and 74%, respectively (Fig. S11). This suggests that when bacteria are swimming backward, they are more likely to enter, navigate and successfully exit confined microenvironments.

**Figure 3:**
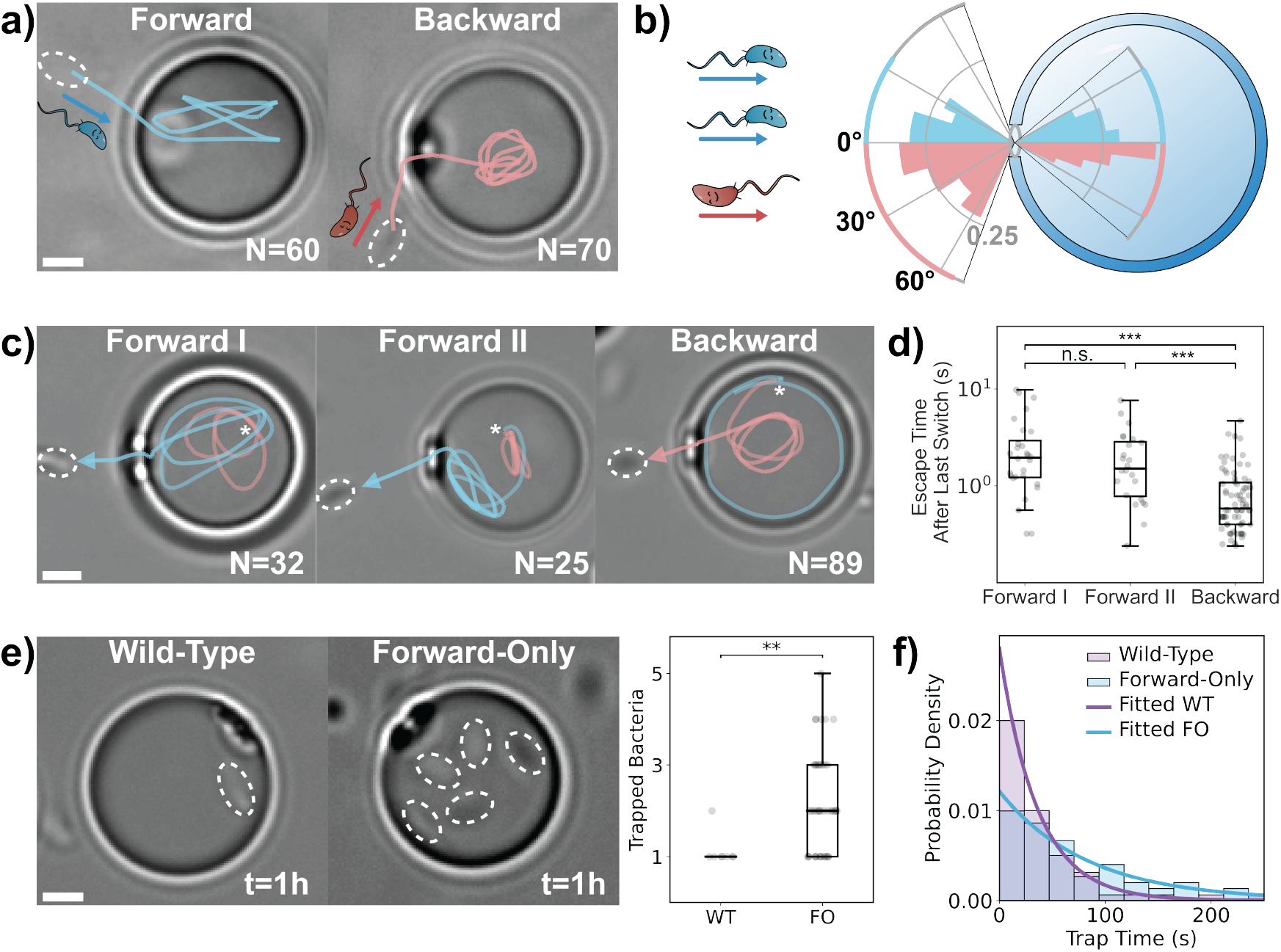
Backward motion enhances entry, exploration, and escape from confined microenvironments. **a)** Representative entry trajectories overlaid on brightfield images. Bacteria approach the µ-Trap either in Forward or Backward mode. N values shown on images; blue – Forward tracks, red – Backward tracks. **b)** Left: cartoon illustrating the Forward–Backward swimming bias outside the trap (∼34% Backward swimmers, N = 182). Right: polar histograms of entry and post-entry angles. Colored arcs denote angular range. Blue: Forward entry/post-entry (N = 28); red: Backward entry/post-entry (N = 43). **c)** Representative escape trajectories overlaid on brightfield images. Inside the µ-Trap, cells exhibit Forward I, Forward II, or Backward motion. N values shown on images; blue - Forward I/II, red - Backward. White asterisks indicate direction-switching events. **d)**Comparison of escape times after the last directional switch. N = 28 for Forward I, N = 24 for Forward II, N = 74 for Backward. ***: *p <* 0.001; Forward I vs. Forward II, *p* = 0.419, not significant (n.s.). **e)** Left: Brightfield images comparing trapping of Wild-Type (WT) and Forward-Only (FO) strains, 1 hour post addition. Right: quantification of the number of trapped cells per filled µ-Trap. *p* = 0.006, **. N = 8 for WT, N = 42 for FO. **f)** Trap residence times fitted to exponential distribution. N = 68 for WT, N = 64 for FO. Dashed white circles in images: bacterial cell bodies. Scale bars: 2 µm.

**Figure 4:**
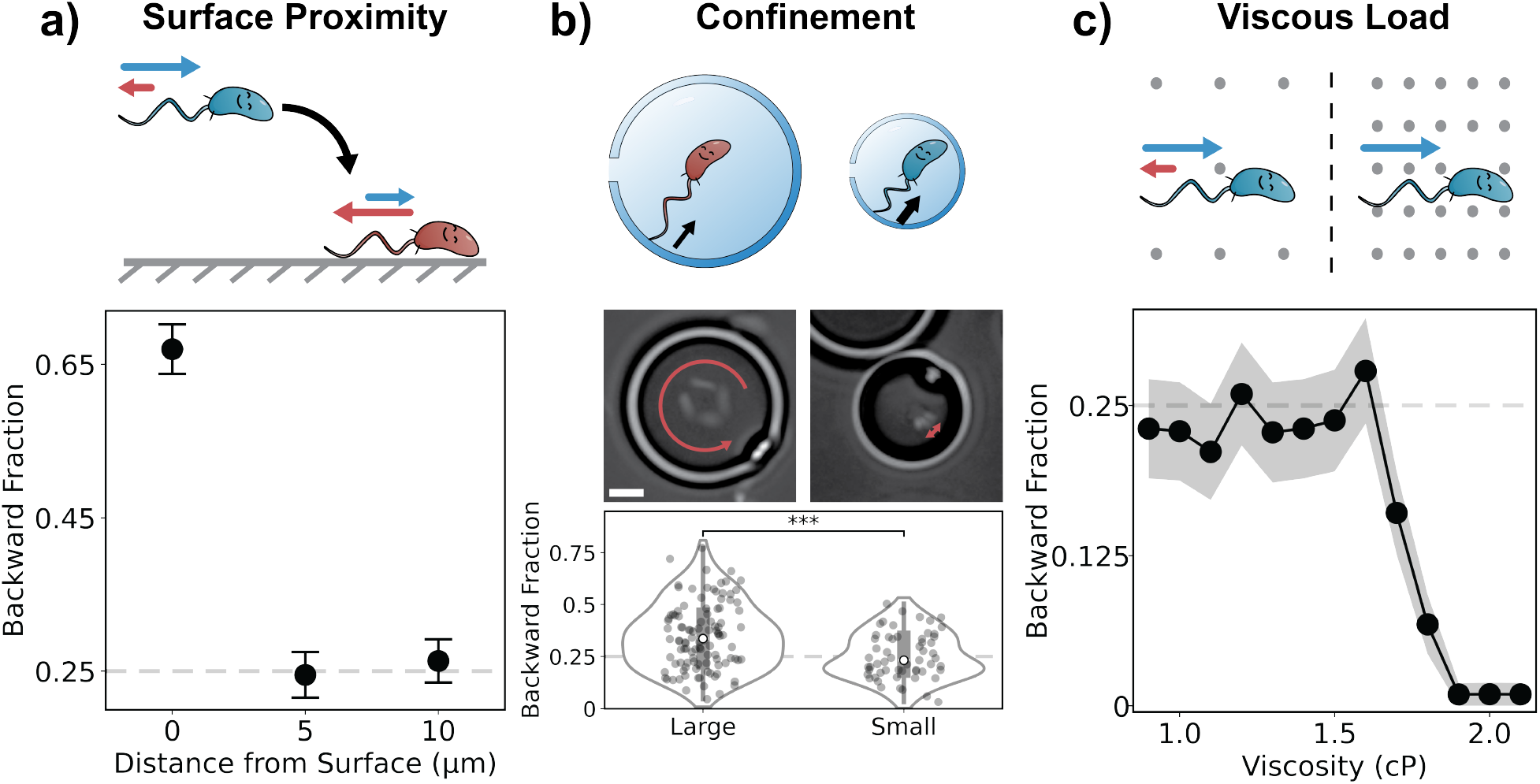
Mechanical load modulates flagellar reversal through a gated switching response. In all cartoons, blue and red bacteria indicate forward- and backward-dominant behavior, respectively; arrow lengths reflect the proportion of each swimming mode. **a)** Cartoons show swimming near surfaces: backward motility is enhanced near the substrate and declines sharply beyond 5 µm. Plot shows the backward-swimming fraction vs. distance from surface. Error bars: standard error of the proportion; N = 208–243 cells per condition. **b)** Cartoons show motility in large and small µ-Traps, with black arrows indicating mechanical load on the flagellum (thin = low, thick = high). Time-lapse overlays (ωt = 0.168 s) show clear backward displacement in large traps (left) and minimal movement in small traps (right); red arcs highlight displacement during backward swimming. Scale bar: 2 µm. Violin plots quantify backward-swimming fractions: 0.35 ± 0.16 in large µ-Traps (N = 108) and 0.26 ± 0.11 in small µ-Traps (N = 57); medians shown as white circles; ***: *p <* 0.001. **c)** Cartoons show response to increasing viscous load. At low viscosity, motility remains reversible; at high viscosity, only forward swimming persists, with gray background dots indicating increased drag. Background gray dots indicate higher drag. Plot shows backward-swimming fraction remains stable up to 1.6 cP, then drops sharply and is abolished above 1.9 cP. N = 101-112 cells per condition. Dashed horizontal lines in plots mark the reference backward-swimming fraction (∼0.25) in unconfined bulk fluid.

To investigate the underlying mechanism, we compared the approach angles of forward and backward entries (Fig. 3b, Fig. S12). Backward swimmers entered over a wider range of angles, with a maximum angle of ∼60°, compared to ∼30° for forward entries, increasing the likelihood of successful navigation. Interestingly, both forward and backward swimmers exhibited similar post-entry maximum angles (∼30°), which also corresponds to the forward maximum entry angle. This suggests that the geometry of the trap and the physical dimensions of the cell body impose a common constraint on alignment after entry. Flagellar labeling further revealed that during backward motion, the flagellum extends ahead of the cell body, facilitating easier exploration into narrow apertures (movie S4).

Escape dynamics revealed another advantage of reverse motion: backward swimmers exited the trap more rapidly than those moving forward (Fig. 3d, movie S5). Although the duration of backward mode was shorter, this alone did not explain the reduced escape time (Fig. S13). This finding aligns with our motion mode characterization (Fig. S8, movie S2), which showed that backward swimmers—despite their slower speeds—exhibit smaller track radii and reduced radial offsets. These features enhance spatial coverage and directional control, suggesting that backward motion enables more efficient navigation in confined environments.

To directly test whether this backward advantage improves exploratory behavior, we compared the Wild-Type (WT) strain with the Forward-Only (FO) mutant. After 1 hour, FO cells exhibited significantly greater trap filling (Fig. S14) and a higher number of cells per trap (Fig. 3e, movie S6). FO swimmers also took more than twice as long to escape (mean escape time: 82.3 s vs. 35.7 s for WT). Fitting these times to exponential distributions yielded Poisson escape rates of *λ*_*WT*_ = 0.0286 s^−1^ and *λ*_*FO*_ = 0.0127 s^−1^ (Fig. 3f). Thus, the ability to reverse doubles the efficiency of escape, highlighting backward motion as a critical adaptive strategy for confined exploration.

Together, these data highlight the pivotal role of backward motion in navigating confined geometries. Rather than being a counter-productive swimming mode, backward motility proves highly efficient for exploring spatially restricted environments. These findings underscore how flagellar reversals function as a specialized and adaptive strategy for maneuvering through complex, three-dimensional spaces.

### Mechanical load gates backward motility across spatial contexts

Previous studies of *Caulobacter crescentus* motility showed that cells spend more time swimming forward than backward (*17,18*). However, these observations span vastly different physical contexts, from unbounded motion in bulk fluid to tethered assays on flat surfaces, raising the question: What physical cues regulate the use of backward swimming?

To isolate the effect of surface proximity, we first quantified the backward fraction of free-swimming cells at defined distances from a substrate. The backward fraction, also referred to as counterclockwise bias (*18*), is the proportion of time a bacterium spends swimming backward relative to its total swimming time. It can be measured from single-cell trajectories or approximated by the fraction of backward-swimming cells in large populations (see methods). When swimming more than 5 µm above the surface, *Caulobacter* cells exhibited a backward fraction of ∼0.25 (Fig. 4a), consistent with previous measurements showing forward-dominant motility in unconfined conditions (*17*). In striking contrast, cells close to the surface showed backward-dominant behavior, with a backward fraction of ∼0.67 (Fig. 4a, movie S7), in agreement with earlier qualitative mobservations of extended backward runs near surfaces (*30*). Our results indicate that surface contact actively promotes backward swimming, likely due to asymmetric mechanical stress experienced during reverse motion near solid boundaries.

Next, we test whether geometric confinement modulates this behavior by tracking cells in colloidal µ-traps of different sizes. In large traps (9.65 ± 1.14 µm in diameter, Fig. S4c), backward motility was enhanced (backward fraction ∼0.35 ± 0.16, Fig. 4b), higher than that of free-swimming cells in open space.. In smaller traps (5.66 ± 0.77 µm in diameter, Fig. S15a), however, the backward fraction dropped to ∼0.26 ± 0.11 (Fig. 4b), due to both shortened backward durations and prolonged forward runs (Fig. S15). Time-lapse overlays and fluorescence imaging confirmed that backward motion produced negligible displacement in small traps, with the flagellum visibly bent against the compartment interior (Fig. 4b, Fig. S16, movie S8). By contrast, forward swimming remained largely unimpeded, with flagella trailing freely. These findings suggest that backward motility operates within a finite mechanical window: moderate force promotes reversals, but excessive load suppresses them. This mechanistic insight may help reconcile previous observations: while Liu *et al*. reported forward-dominant motility under minimal mechanical load in bulk fluid (*17*), and Lele *et al*. observed further suppression of backward motion under high-load tethered conditions (*18*), our system bridges both extremes within a single geometric continuum. Notably, we identified an intermediate regime where cells actively leverage their highly efficient backward motility to navigate their environment.

To further isolate the effect of mechanical load symmetry, we compared µ-Trap confinement with viscous drag. In tight traps, mechanical load is asymmetric—backward swimming presses the flagellum against the boundary, while forward swimming primarily exerts force on the cell body. Conversely, high-viscosity media distribute mechanical load symmetrically along the body and filament. This distinction, highlighted by the contrast between Fig. 4b and Fig. 4c, suggests that directional switching is more sensitive to localized asymmetric stress than to uniform drag. To systematically test this force-dependent gating, we titrated medium viscosity using glycerol (Fig. S17). At low to moderate viscosities (0.9–1.6 cP), the backward fraction remained near 0.25 (Fig. 4c). However, beyond 1.6 cP, backward motion sharply declined and was abolished above 1.9 cP (movie S9), consistent with a threshold-like response to mechanical load. Notably, while surface contact enhanced backward motility, elevated viscosity did not (Fig. 4a vs. 4c), reinforcing the idea that *Caulobacter* distinguishes between asymmetric mechanical stresses and symmetric viscous forces.

To support this distinction, live cell imaging showed that contact between the flagellum and the surface µ-Trap triggered an immediate directional switch, while collisions of the cell body did not (Fig. S18, movie S10). These observations imply that mechanical stress localized to the flagellum, not the body, regulates motor switching, likely by modulating the dissociation of response regulators such as the CheY-P (phosphorylated CheY) and Cle proteins from the switch complex (*10,15,16,31*).

Together, these findings establish that the *Caulobacter crescentus* flagellar motor functions as a gated mechanosensor, translating local mechanical inputs into adaptive changes in swimming behavior. In open fluid, cells maintain a low backward fraction to balance propulsion and exploration. Near surfaces or under moderate confinement, backward swimming is selectively enhanced to aid in reorientation and escape. Under high load—such as in tight confinement or viscous environments—backward motility is actively suppressed, conserving energy and avoiding ineffective motion. Through this load-sensitive tuning of motor activity, *Caulobacter* dynamically tailors its motility strategy to the mechanical landscape of its three-dimensional surroundings.

## Discussion

Our findings reframe backward swimming not as a redundant behavior but as a context-sensitive strategy optimized for physical confinement. Using ecological assays and precisely engineered colloidal µ-Traps, we show that flagellar reversibility enables *Caulobacter* to access nutrient-rich niches inaccessible to mutants lacking backward motility. Within µ-Traps, backward motion aligns the cell body and flagellum for efficient entry and escape through narrow apertures—geometrically favorable maneuvers that forward swimming alone cannot achieve.

Critically, we reveal that reversals are not stochastic but triggered by a finely tuned mechanical switch. Moderate mechanical stresses, such as surface contact, stimulate reversals, likely via conformational changes that modulate CheY-P and Cle protein interactions at the motor. In contrast, high-load conditions such as tight confinement or elevated viscosity, suppress reversal, effectively locking the motor into a single direction. This gating mechanism ensures that reversals are selectively deployed when advantageous and suppressed when energetically costly or ineffective.

Together, these findings offer two interconnected but distinct insights: backward motility is strategically deployed to overcome physical confinement (*why*), and this behavior is regulated by a force-sensitive molecular mechanism at the flagellar motor (*how*). While each insight stands on its own, it is their coupling: precisely tuned reversal in response to mechanical constraints, that enables efficient bacterial navigation in complex 3D environments. These principles may extend to other *monotrichous* swimmers, including pathogens such as *Pseudomonas aeruginosa* and *Vibrio cholerae*, which encounter complex spatial constraints in host environments. Our results suggest that physical parameters like viscosity and confinement geometry can directly regulate intracellular signaling states, offering new levers for manipulating microbial behavior.

By enabling long-term observation of free-swimming bacteria under realistic confinement, the µ-Trap system overcomes limitations of conventional 2D assays and unstructured 3D chambers (*17, 18*). This architecture challenges classical run-and-tumble paradigms derived from *E. coli* and uncovers finely tuned reversal behavior in *monotrichous* species like *Caulobacter*, bridging a longstanding gap between theoretical models of confined locomotion and direct experimental observation.

Despite its power, the µ-Trap platform has limitations. Current designs do not permit simultaneous imaging of intracellular structures, and adapting trap geometries for different species requires fabrication redesign, though emerging reconfigurable traps offer promising solutions (*29*). Future advances integrating chemical gradients and surface chemistries will further expand the platform’s capabilities.

Together, the µ-Trap approach, coupled with 3D imaging and real-time tracking, establishes a versatile system to dissect bacterial navigation strategies under realistic physical constraints. More broadly, our results show how bacteria convert mechanical stress into adaptive motile behavior—revealing a mechanical logic for navigating life’s tight spots.

## Acknowledgments

We would like to thank the members of the Sacanna Lab and the Saurabh Lab for their invaluable support. Our sincere gratitude also goes to the Brun Lab, the Jenal lab, and the Thanbichler Lab for generously sharing strains and plasmids. Special thanks to M. Panjalingam and A. Chimthanawala from the Saurabh Lab for their assistance with strain construction.

## Author contributions

S.S.1, S.S.2, and R.Z. conceived the study. R.Z. performed experiments with inputs from S.S.1 and S.S.2. R.Z., analyzed the data. R.Z. and S.S.2 developed image processing and tracking tools. S.S.1, S.S.2 and R.Z. wrote the manuscript.

## Data and materials availability

All data supporting the findings of this study are available in the main text and supplementary materials. Raw image datasets and analysis code are available at: https://github.com/saurabhLabNYU/Caulobacter_Reversal_Gating.

## Supplementary materials

Materials and Methods

Figures S1 to S18

Table S1

References (*32-43*)

Movies S1 to S10

