## Supplementary Information for "A Mechanical Logic for Bacterial Navigation"

**This PDF file includes:**

Materials and Methods

Figures S1 to S18

Table S1

Captions for Movies S1 to S10

**Other Supplementary Materials for this manuscript:**

Movies S1 to S10

### Materials and Methods

The bacterial strains, plasmids, and other resources used in this study are listed in **Table S1**. This includes detailed information on the strains and plasmids, as well as additional materials such as chemicals, reagents, and software tools employed throughout the study.

### Bacterial Cell Growth

*Caulobacter crescentus* strains were grown in Peptone Yeast Extract (PYE) medium (*32*) at 30°C with shaking at 200 rpm overnight. No additional antibiotics were required for the strains used in this study. Prior to experiments, overnight cultures were diluted into M2G medium (M2 medium supplemented with 0.2% w/v glucose) (*33*) and grown to mid-log phase.

Swimmer cells were isolated using established synchronization protocols (*34*). Following synchronization, cells were resuspended in M2 medium (without glucose) and incubated at 30°C for 10 minutes to recover from cold shock induced during the synchronization step. The optical density at 600 nm ($OD_{600}$) was measured using a UV-Vis spectrophotometer, and the culture was diluted with M2 medium to a final $OD_{600}$ of 1.0.

To characterize swimmer cell length, a 1 µL aliquot of the prepared culture was added on a coverslip and sealed under a 1.5 wt% M2 agarose pad. Cell lengths were quantified using MicrobeJ.

### Filter Paper Experiment Setup

In a typical experiment, filter paper (Grade A-E glass microfiber filters, 1.0 µm particle retention) was cut into approximately 2 cm × 2 cm squares. Prior to use, the filter papers, Petri dish, and tweezers were sterilized under UV light for 2 hours.

A 1.5 wt% agarose solution was prepared using PYE medium and poured into a Petri dish. A piece of cut filter paper was placed on the agarose surface and allowed to hydrate for 5 minutes.

Wild-Type (NA1000) and Forward-Only mutant (NA1000 Δ*cheR1* Δ*cheR2* Δ*cheR3*) swimmer cells were prepared as described above and diluted with M2 medium to $OD_{600}$ = 0.05. Then, 10 µL of the diluted cell suspension was added onto the wetted filter paper. After 3 minutes, the filter paper was removed, and the agarose plate was incubated at 30°C for 40 hours.

Images of resulting colonies were acquired using a smartphone camera and processed in FIJI. Colony-forming units were counted. Each condition was tested with 6 biological replicates.

As controls, 10 µL of M2 medium (without cells) was added onto filter paper for the negative control, and 10 µL of diluted swimmers was added directly onto the agarose without filter paper for the positive control.

To compare the survival ability of the two bacterial strains, a serial dilution assay was performed. 10 µL of swimmer cultures diluted in M2 medium (undiluted, $10^{-3}$, $10^{-4}$, $10^{-5}$, $10^{-6}$, and $10^{-7}$) were directly spotted onto agarose plates.

The geometry and mesh structure of the filter paper were characterized using scanning electron microscopy (SEM). Small sections of the filter paper were cut and mounted directly onto conductive tape without additional coating. Imaging was performed at an accelerating voltage of 3 kV.

### Synthesis and Characterization of µ-Traps

The synthesis of µ-Traps was based on a previously reported method (*29*) with slight modifications to accommodate the size of bacteria(Fig. S3).

3-(Trimethoxysilyl)propyl methacrylate (TPM) oil was added to deionized water at a 1:10 volume ratio and hydrolyzed under gentle rotation at 30 rpm until the solution became clear. A total of 7.5 mL of this hydrolyzed TPM solution was mixed with 100 µL 16.3 wt% 1.3 µm negatively charged polystyrene (PS) spheres, 112.5 µL 100 mM NaOH, and deionized water to bring the final volume to 15 mL. The mixture was transferred to a centrifuge tube mounted on a Roto-Mini Plus (Ward's Science) rotating at 10 rpm to prevent sedimentation.

After 23 hours, the resulting TPM emulsion with PS heads was added to a NaOH solution (final $[OH^-]$ = 100 mM) to induce a self-inflation process. The inflation proceeded for 20 minutes, with the emulsion rotated once every 5 minutes. Polymerization was initiated by adding 0.1 vol% of the photo-initiator Darocur® 1173, followed by UV irradiation for 25 minutes.

The polymerized particles were washed in water and then transferred to tetrahydrofuran for further cleaning to remove the PS heads. The particles were subsequently washed back into water. Each washing step consisted of three rounds.

To produce small µ-Traps, the inflation time was reduced to 1 minute. However, unless otherwise specified, large µ-Traps were used in most experiments throughout this study.

The optical density of the prepared µ-Trap suspension was measured at 450 nm using a UV-Vis spectrophotometer.

The overall diameter of the µ-Traps was measured using brightfield imaging, as detailed in the following section. Diameter measurements were extracted using a custom Python script based on the Hough transform.

The diameter of the µ-Trap apertures was characterized by SEM using a 2.5 nm platinum coating and a 1 kV accelerating voltage. Aperture sizes were quantified with the same Hough-transform-based Python analysis.

To enhance the 3D visualization of the µ-Traps in Fig. 2b, the SEM stage was tilted at an angle of 40° during imaging.

### Brightfield Microscopy Setup

Imaging was performed on an inverted Nikon Eclipse Ti2-E microscope equipped with a 100× oil-immersion objective (Plan Apochromat $\lambda$D, N.A. 1.45) at room temperature.

Prior to imaging, a large µ-Trap stock solution was diluted in M2 medium and stabilized with the surfactant Pluronic® F108 (hereafter referred to as F108). The dilution was prepared by sequentially mixing 190 µL M2, 4 µL of 0.5 wt% F108 in deionized water, and 6 µL of large µ-Trap stock ($OD_{450}$ = 3.5). In a 384-well glass-bottom plate, 67.5 µL M2 and 18 µL of the diluted µ-Trap solution were added and allowed to sediment for at least 20 minutes. Then, 4.5 µL of prepared swimmer cells ($OD_{600}$ = 1.0) was added. The total volume per well was 90 µL, with final concentrations of 0.002 wt% F108, µ-Trap $OD_{450}$ = 0.021, and swimmer $OD_{600}$ = 0.05.

To visualize and analyze the motion of Caulobacter crescentus swimmer cells within the µ-Traps, images were acquired under brightfield illumination at 47 frames per second (FPS) with a 1024 × 1024 pixel field of view. No dichroic mirrors or optical filters were used. Each video typically lasted 3 minutes.

For experiments comparing large and small µ-Traps, a mixed solution was prepared by combining 187 µL M2, 4 µL of 0.5 wt% F108, 3 µL of small µ-Traps ($OD_{450}$ = 7.0), and 6 µL of large µ-Traps ($OD_{450}$ = 3.5). A total of 67.5 µL M2 and 18 µL of this mixed solution were added to a well, followed by 4.5 µL of swimmer cells. The final concentrations were 0.002 wt% F108, $OD_{450}$ = 0.021 for each µ-Trap size, and $OD_{600}$ = 0.05 for swimmer cells.

To facilitate the detection of entry and escape events, imaging was performed at a larger field of view (1416 × 1416 pixels) at a reduced frame rate of 20 FPS. Videos of 12 minutes in duration were typically acquired under these conditions.

The motile fraction of swimmer cells was observed to decrease around 20 minutes after addition. Therefore, each well was imaged continuously for no more than 12 minutes post addition.

However, to compare trapping behavior between Wild-Type and Forward-Only strains, videos were acquired at 20 FPS one hour after swimmer cells addition.

### Effect of Surfactant on Bacterial Motility

To assess the effect of surfactant on bacterial motility, brightfield imaging was performed using the same optical setup as described above. A diluted F108 solution was prepared by mixing 196 µL M2 medium with 4 µL of 0.5 wt% F108 in deionized water.

In a 384-well glass-bottom plate, two conditions were prepared: (1) 67.5 µL M2 + 18 µL F108 solution + 4.5 µL of prepared swimmer culture, resulting in a final F108 concentration of 0.002 wt%; (2) 85.5 µL M2 + 4.5 µL of prepared swimmer culture, with no added surfactant (final F108 = 0 wt%). The final volume in both conditions was 90 µL, and the swimmer cell density was adjusted to $OD_{600}$ = 0.05.

Videos were recorded at 30 FPS. Bacterial trajectories were analyzed using TrackMate in FIJI and classified using Ilastik. The median speed of each trajectory was computed and compared between conditions.

### 3-Dimensional Tracking Setup

It was observed that the apparent shape of bacterial cells varied with their axial (z) position under brightfield imaging. To establish a quantitative relationship between the z-coordinate and bacterial shape, a 3D reference library was created.

Some swimmer cells naturally adhered to the glass surface of a 384-well plate in M2 medium. Z-stacks of these adhered cells were acquired using a step size of 0.1 µm. For each z-slice, the bacterial shape was segmented via thresholding and fitted with an ellipse using a custom Python script. A consistent correlation was identified between the z-position and the minor axis length of the fitted ellipse. Since this relationship was sensitive to the choice of thresholding and lacked a straightforward analytical form (Fig. S6), it was empirically modeled using two separate single-exponential functions—one for $z > 0$ and another for $z < 0$—assuming $z = 0$ at the focal plane(Fig. S5a).

This empirical shape-to-depth relationship was further validated by comparison with simulated point spread functions (PSF), generated using a previously reported brightfield PSF model (*35*). Simulations were performed using a wavelength of 450 nm, a numerical aperture (N.A.) of 1.45, and a refractive index n = 1.518 for immersion oil.

To quantify the robustness of this calibration, a dataset of 75 flat swimmer cells was split into training (N = 60) and testing (N = 15) sets. The single-exponential model was fit to the training set, and prediction error was evaluated across the training, testing, and an additional validation set comprising 30 tilted swimmer cells that adhered at an angle (Fig. S5b).

For 3D tracking within µ-Traps, 2D trajectories of both swimmer cells and µ-Traps were first identified using TrackMate in FIJI and Ilastik. The positional data were then passed to a custom Python pipeline that extracted the bacterial shape and estimated 3D coordinates using the calibrated ellipse-minor-axis fitting approach. The same image processing pipeline—including erosion and filtering steps—was applied to remove transient passersby and background noise, in accordance with the procedures used during model fitting.

Finally, the 3D trajectories were visualized using custom Python code, and the motion of the µ-Trap was subtracted to yield swimmer trajectories normalized relative to the µ-Trap.

### Flagellar Labeling and Fluorescence Microscopy

To visualize flagella, a synchronizable strain (NA1000 *fljK::T103C*) was constructed as previously described (*36*). Swimmer cells were isolated as described above.

For flagellar labeling, a 5 mM Alexa Fluor$^{TM}$ 546 (AF546)-maleimide dye stock was prepared in dimethyl sulfoxide (DMSO). A total of 198 µL of synchronized swimmer cells ($OD_{600}$ = 1.0) was mixed with 2 µL of the dye to achieve a final concentration of 50 µM, while keeping DMSO below 5 vol%. The mixture was incubated at 30°C for 30 minutes and then washed twice with 1 mL M2 medium (centrifugation: 15000 rcf, 2 minutes). Cells were resuspended in 200 µL M2 medium.

Alexa Fluor™ 488 (AF488)-maleimide dye was tested under the same conditions using the same final dye concentration.

To assess flagellar labeling, the stained culture was diluted 8-fold. 1 µL of the diluted culture was spotted onto a glass coverslip and sealed under a 1.5 wt% agarose pad prepared with M2 medium.

For dual labeling of the flagella and cell body, 200 µL of AF546-labeled culture was supplemented with 1 µL of 500 µM Nile Blue A in DMSO, yielding a final Nile Blue A concentration of 5 µM. No washing was performed after Nile Blue A addition.

It was observed that swimmer motility decreased after ∼1 minute of continuous exposure to 488 nm TIRF laser. To quantify this effect, unlabeled Wild-Type swimmers were imaged under 488 nm or 561 nm lasers (5–100% intensity) along with brightfield illumination. Laser power density was measured using a power meter (ThorLabs PM160) (561 nm: $3.9 \times 10^3$–$4.1 \times 10^4$ mW/cm$^2$; 488 nm: $2.3 \times 10^3$–$3.1 \times 10^4$ mW/cm$^2$). 2D trajectories were analyzed using TrackMate in FIJI and Ilastik, and median speed was compared at different timepoints. Due to the additional laser exposure, 3D tracking was not applied in these conditions.

To quantify Brownian motion and compare it with decreased motility, hydrogen peroxide ($H_2O_2$) was added to final concentrations of 0.01–10 wt%. No active swimming was observed at or above 0.01 wt% $H_2O_2$. For speed analysis, cells exposed to 5 wt% $H_2O_2$ were tracked using the same pipeline.

To image AF546-labeled flagella in µ-Traps, samples were illuminated with 10% TIRF 561 nm laser ($6.5 \times 10^3$ mW/cm$^2$) and 10% brightfield illumination. Exposure time was 30 ms. The 561 nm laser excited AF546-labeled flagella, while brightfield illumination revealed the swimmer body and µ-Trap geometry. A 607/36 nm emission filter was used.

For AF488-labeled flagella, imaging was performed using 10% TIRF 488 nm laser ($4.1 \times 10^3$ mW/cm$^2$) and 25% brightfield illumination with a 30 ms exposure. A 525/30 nm emission filter was used.

For dual-label imaging of AF546-labeled flagella and Nile Blue A-stained cell bodies, two-channel fluorescence imaging was performed using 10% TIRF 561 nm laser. A dichroic mirror

(ZT640rdc-xr-UF2) and two emission filters (607/36 nm and 700/75 nm) were used. Exposure time was 30 ms.

### Analysis of µ-Trap Videos

For a typical analysis pipeline, regions containing trapped swimmers were manually labeled and cropped using FIJI.

**Motion Mode Characterization**

Three distinct motion modes were identified: Forward I, Forward II, and Backward. Each video was manually annotated with the corresponding motion mode and segmented accordingly. 3D trajectories from unique swimmers were analyzed as described previously. Key metrics—including 3D speed, distance from surface, circular trajectory radius, and radial offset—were calculated and compared using custom Python code. The dataset included 34 Forward I, 30 Forward II, and 56 Backward trajectories from unique swimmers.

**Entry, Escape, and Trapping Events**

For entry and escape analysis, the timepoints of entry and escape were recorded manually. Entry and post-entry angles were determined based on 3D positions before and after entry (defined as the farthest position before the swimmer changed direction). Of 150 observed entry events, 71 yielded analyzable angles. Events were excluded if the exterior trajectory was untrackable (e.g., swimmer moved out of focus or overlapped with a passerby) or if the trap aperture was not facing sideways.

Escape times were calculated as the duration between the escape timepoint and the most recent Forward-to-Backward or Backward-to-Forward switching event. No transitions between Forward I and Forward II were typically observed during this interval. Among 147 observed escape events, 126 were analyzable (exclusions were made if multiple swimmers were present in a single µ-Trap).

When both entry and escape events were observed for the same swimmer, the trapping time was calculated and fitted with an exponential distribution, assuming a Poisson process for escape. Trap times from 68 Wild-Type and 64 Forward-Only swimmers were analyzed and compared.

Trapping behavior between Wild-Type and Forward-Only strains was further compared by evaluating µ-Trap occupancy. Videos acquired 1 hour post addition were analyzed using a custom Python script to detect µ-Trap counts. The number of filled µ-Traps was manually identified, and the filling percentage was computed. For each condition, data from 4 fields of view (each containing 11–21 µ-Traps) were compared. Additionally, the number of trapped swimmers in each filled µ-Trap was recorded and compared (8 filled µ-Traps for Wild-Type and 42 for Forward-Only).

### Measurement of Backward Fraction

The backward fraction is defined as the proportion of time a swimmer exhibits backward motion, which corresponds to counterclockwise (CCW) rotation of the flagellum (*18*). This is also referred to as the CCW bias.

To compare backward fractions under different confinement conditions, videos of swimmers in fields containing both large and small µ-Traps were acquired as described above (47 FPS). Regions containing trapped swimmers were manually cropped and labeled by motion mode. For calculating the backward fraction, Forward I and Forward II modes (corresponding to clockwise flagellar rotation) were combined. Mode durations at the start and end of each trajectory were excluded. The backward fractions from 108 unique swimmers in large µ-Traps and 57 unique swimmers in small µ-Traps were analyzed.

To quantify confinement differences, the available space (defined as the inner diameter of each µ-Trap) and swimmer body length were measured using FIJI and compared.

When sample sizes were sufficiently large, the backward fraction was also estimated from the number fraction—i.e., the proportion of swimmers exhibiting backward motion at a given time point. It was found that the backward fraction converged to a stable value when the sample size exceeded 100 swimmers. Frames were sampled at 12-second intervals to avoid duplicate counting.

To assess the backward fraction outside of µ-Traps, flagella were labeled with AF546 as described above, and samples were prepared in standard wellplate format containing only large µ-Traps. The backward fraction was measured from 182 freely swimming cells located outside the µ-Traps, and the standard error of the proportion was calculated.

To investigate how proximity to surfaces affects the backward fraction, dual labeling of the cell body (Nile Blue A) and flagella (AF546) was used. Samples were imaged at three focal depths: 0, 5, and 10 µm above the glass surface. Swimmers were counted at each depth (N = 212, 208, and 243, respectively). The shallow depth of field in Fluorescence imaging ensured exclusion of swimmers outside the focal plane.

To assess how mechanical load influences the backward fraction, swimmers with AF546-labeled flagella were suspended in M2 medium containing increasing glycerol concentrations (0–26 vol%) to modulate viscosity. Temperature was maintained at 25°C using an Okolab stage heater. Imaging was performed 5 µm above the surface using 10% TIRF 561 nm laser and 10% brightfield illumination to extend depth of field and minimize out-of-focus tracking. Five timepoints spaced 12 seconds apart were examined. At each timepoint and its adjacent frames, the swimmer's motion direction (forward or backward) was recorded whenever the flagellum was in focus, allowing determination of the motion mode at that timepoint. For each condition, the swimming direction of 101–112 swimmers was determined, and standard error of the proportion was calculated.

To determine the relationship between glycerol concentrations and fluid viscosity, glycerol-M2 mixtures (0, 15.9, and 27.4 vol%) were prepared and measured at 25°C using a glass capillary viscometer. Each measurement was repeated three times. The measured dynamic viscosities were in good agreement with classical values for glycerol–water mixtures (*37*). The experimental glycerol concentrations used corresponded to dynamic viscosities in the range of 0.9–2.1 cP (with a step size of 0.1 cP) at 25°C.

### Statistical Analysis

All statistical analyses were performed using custom Python code. The normality of the datasets was first assessed using the Shapiro-Wilk test. If the data did not follow a normal distribution (Shapiro-Wilk $p < 0.05$), the non-parametric Mann-Whitney U test was applied for pairwise comparisons. If the data followed a normal distribution, a Student's $t$-test was used. Statistical significance is indicated as follows: *** ($p < 0.001$), ** ($p < 0.01$), * ($p < 0.05$), n.s. (not significant). For $p > 0.01$, the exact $p$-value is reported.

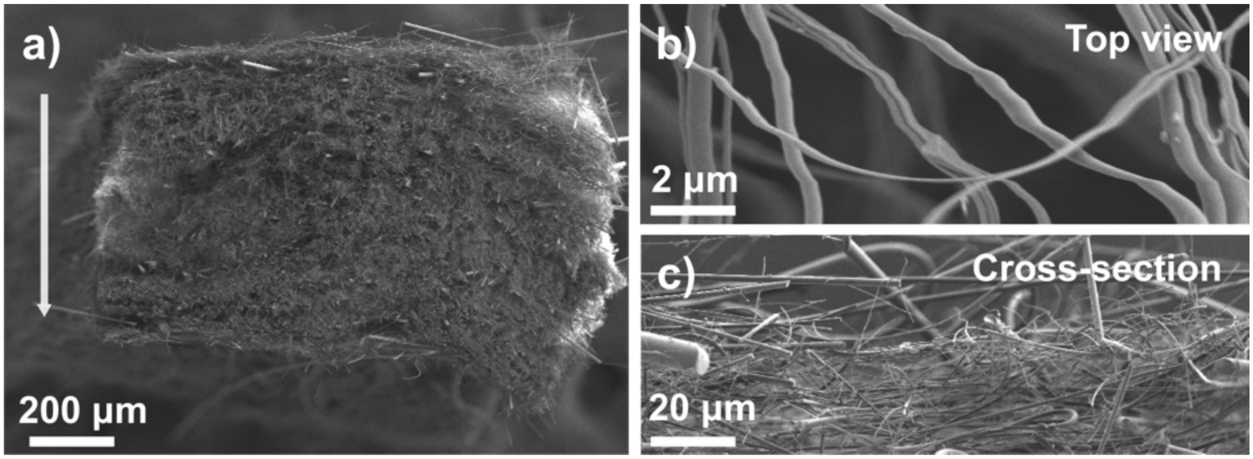


**Figure S1**: **Scanning electron micrographs of filter paper. a)** Cross-sectional view; the arrow indicates the swimming direction of swimmer cells passing the filter. **b)** Top-down view of the filter surface. **c)** Magnified cross-sectional view highlighting the microstructure of the filter paper.

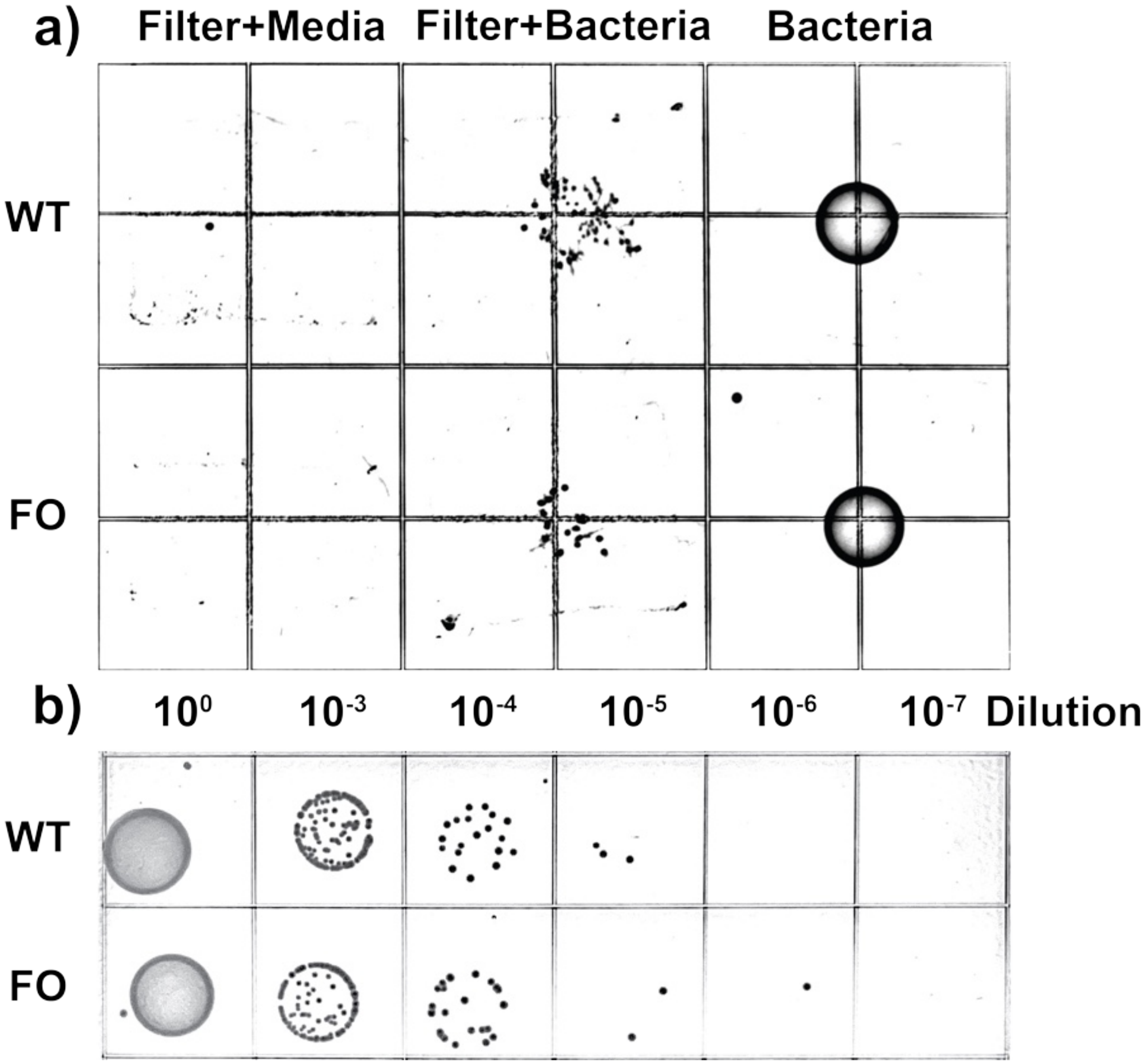


**Figure S2**: **Controls and survivability assay for swimmer cells. a)** Experimental setup: negative control (filter + media), experimental group (filter + bacteria), and positive control (bacteria without filter). **b)** Serial dilution results showing no significant difference in survivability across groups. WT: Wild-Type; FO: Forward-Only.

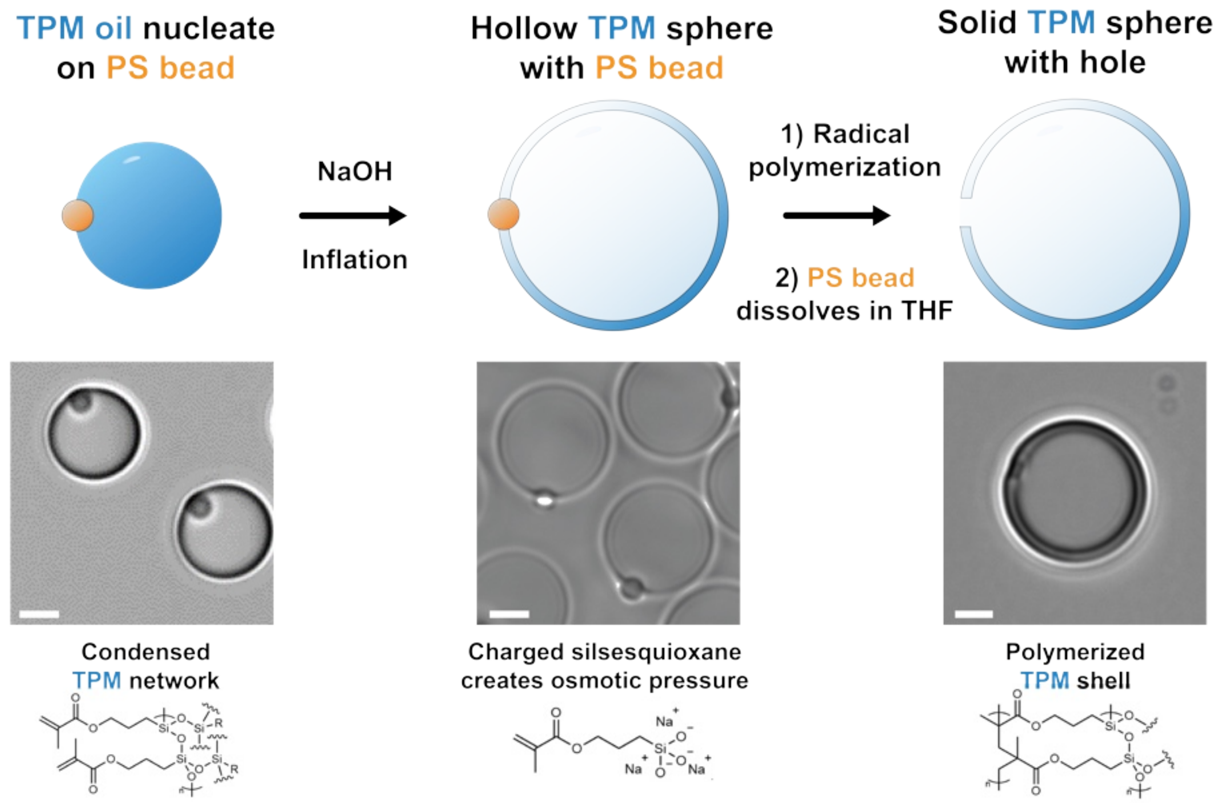


**Figure S3**: **Schematic of µ-Trap synthesis process.** The process includes four main steps: nucleation, inflation, polymerization, and washing.

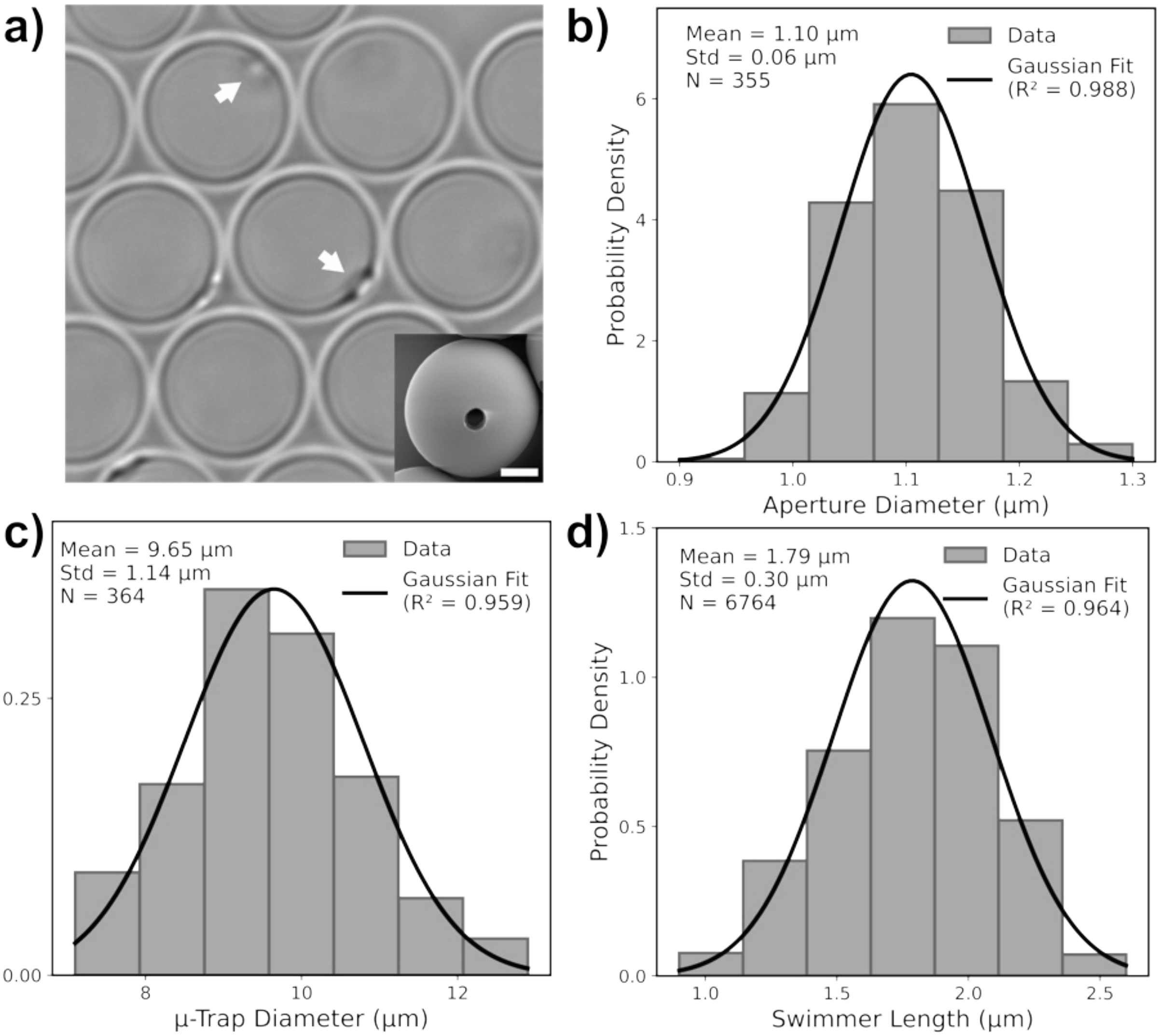


**Figure S4**: **Characterization of synthesized µ-Traps and swimmer cells. a)** Brightfield image of synthesized µ-Traps. Arrows indicate the apertures. Inset: SEM image. Scale bar: 2 µm. **b)** Distribution of aperture diameters (N = 355). **c)** Distribution of µ-Trap body diameters (N = 364). **d)** Distribution of *Caulobacter crescentus* swimmer lengths (N = 6764).

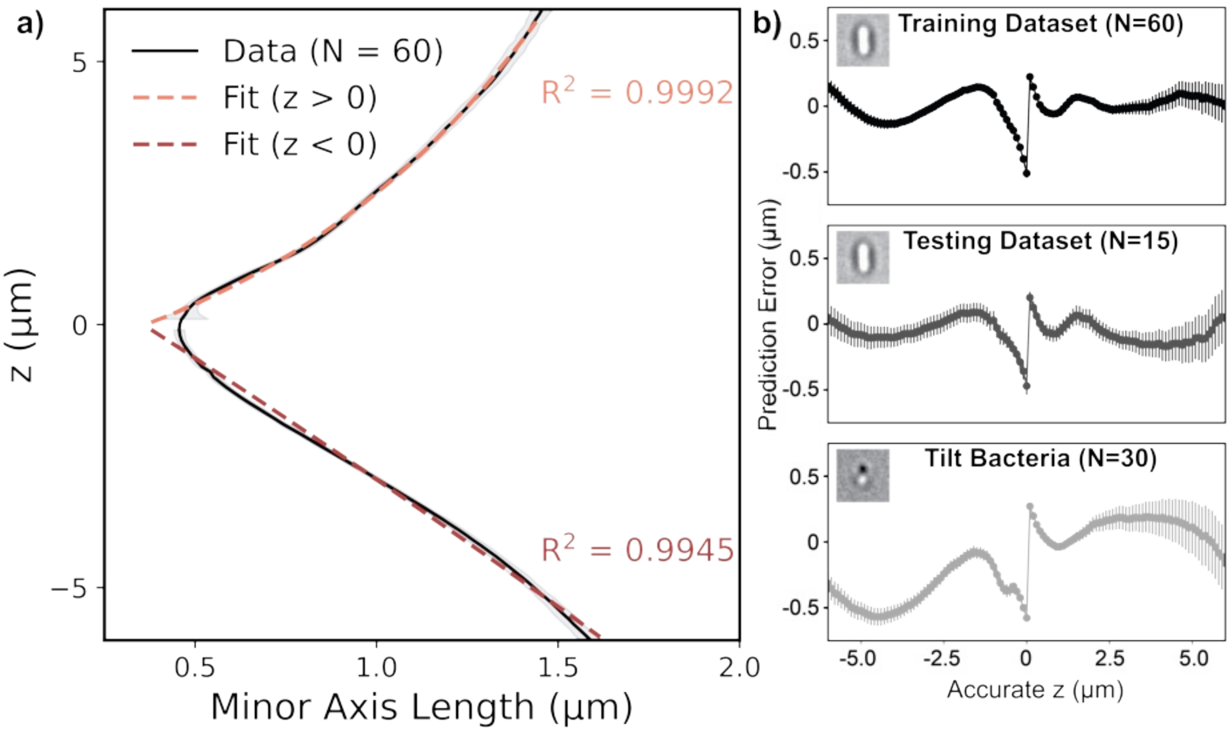


**Figure S5**: **Calibration of z-minor axis relationship and model validation. a)** Relationship between z-position and minor axis length, based on a training dataset of 60 flat Caulobacter swimmer cells. Separate single-exponential fits were applied to z > 0 and z < 0 regions. Shaded areas represent 99% confidence intervals. **b)** Prediction error of the fitted z–minor axis relationship evaluated on the training dataset (60 flat swimmers), a testing dataset (15 flat swimmers), and a tilt dataset (30 swimmers tilted at an angle on the surface).

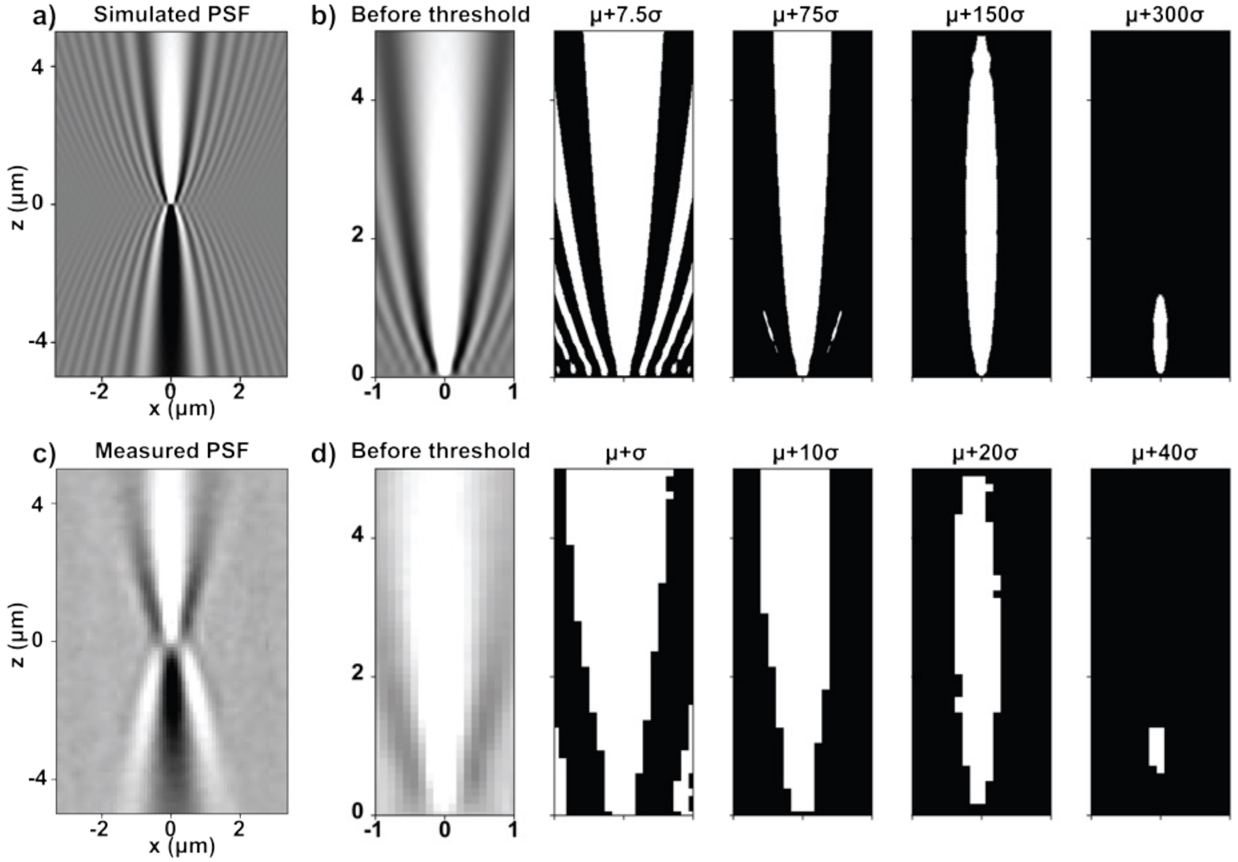


**Figure S6**: **Simulated and experimental point spread function (PSF) and thresholding effects.** **a)** Simulated PSF illustrating the relationship between z-position and minor axis length. **b)** The z–minor axis relationship is influenced by the thresholding process. Threshold = $\mu$ + n$\sigma$, where $\mu$ is the background mean and $\sigma$ is the background standard deviation. **c)** Experimentally measured PSF of a flat, surface-adhered bacterial swimmer. **d)** Resulting PSF outlines after applying different thresholding levels.

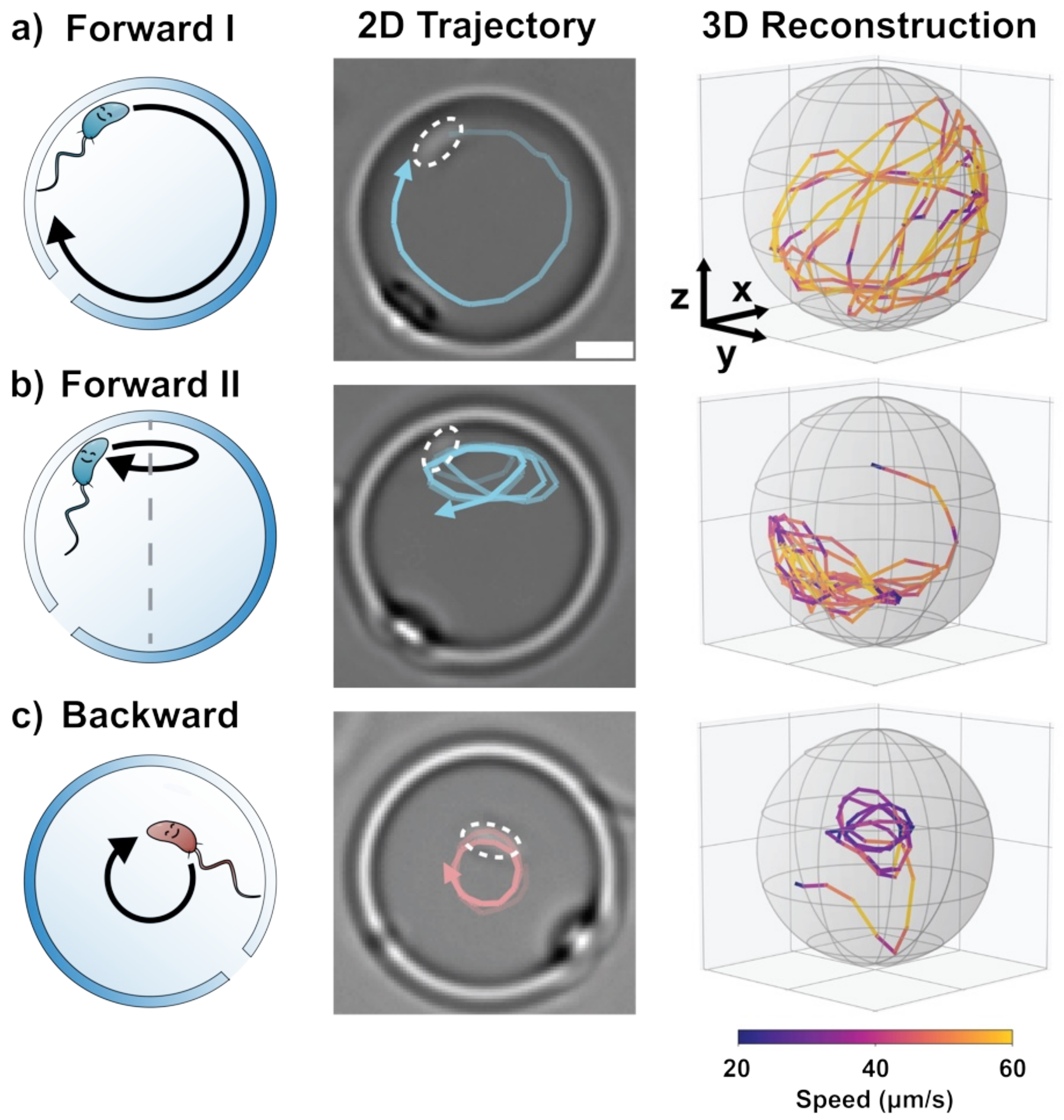


**Figure S7**: **Characterization of swimmer motion modes in μ-Traps.** From left to right: schematic illustration, 2D trajectory overlaid on brightfield images, and 3D trajectory reconstruction. **a)** Forward I motion. **b)** Forward II motion. **c)** Backward motion. Blue tracks represent forward motion; red tracks represent backward motion. Scale bar: 2 μm.

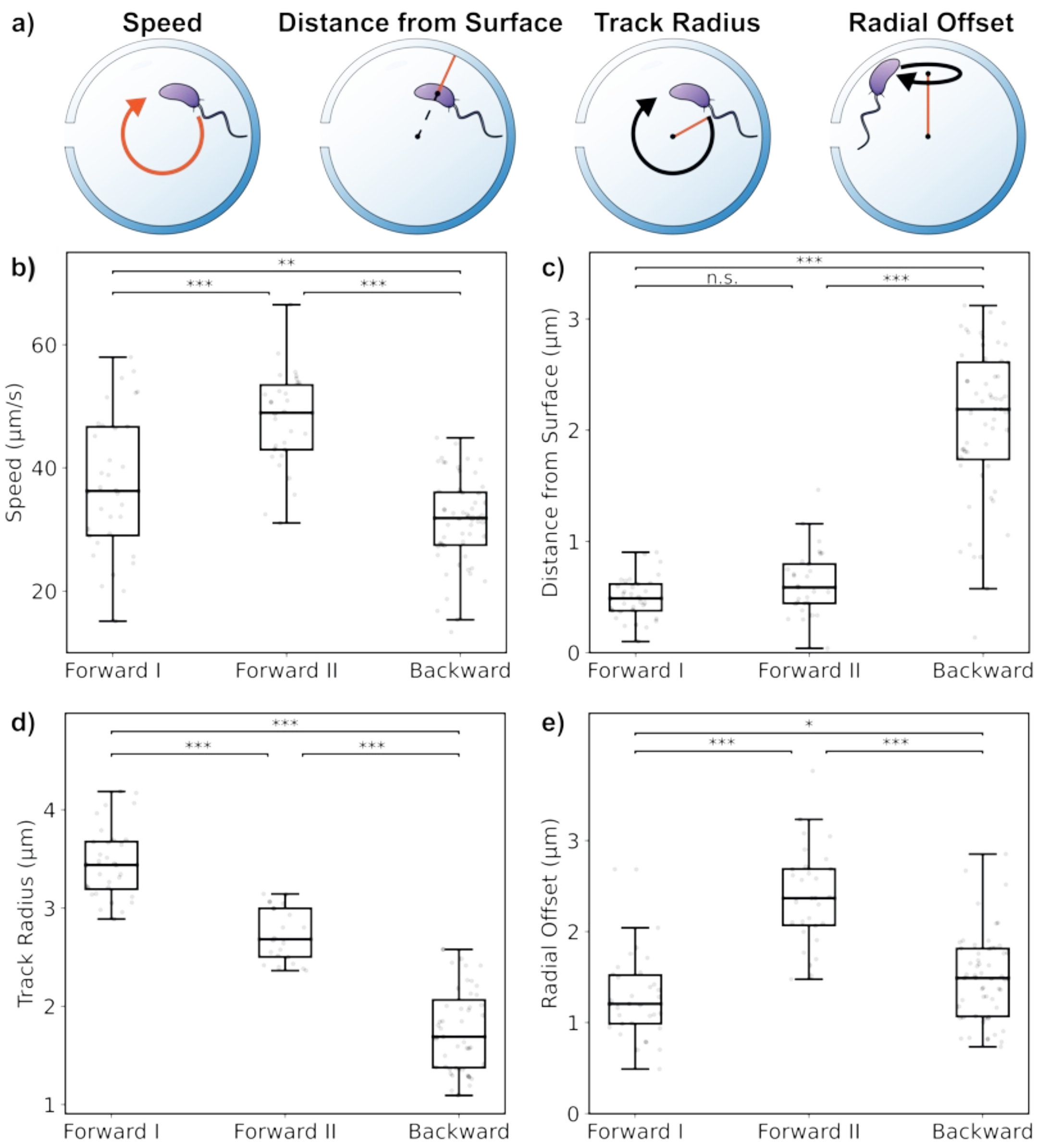


**Figure S8**: **Comparison of motion modes.** (N = 34 for Forward I, N = 30 for Forward II, N = 56 for Backward; ***: $p < 0.001$; **: $p < 0.01$; *: $p < 0.05$; n.s.: not significant. $p$-values greater than 0.001 are indicated numerically.) **a)** Illustration of the key parameters compared. Speed: 3D speed of the bacterial swimmer. Distance from surface: distance between the center of mass of the swimmer and the inner surface of the µ-Trap. Track radius: radius of the circular trajectory followed by the swimmer. Radial offset: distance between the center of the circular trajectory and the center of the µ-Trap. **b)** Speed comparison (Forward I vs. Backward, $p = 0.00972$, **). **c)** Distance from surface comparison (Forward I vs. Forward II, $p = 0.0568$, n.s.). **d)** Track radius comparison. **e)** Radial offset comparison (Forward I vs. Backward, $p = 0.0394$, *).

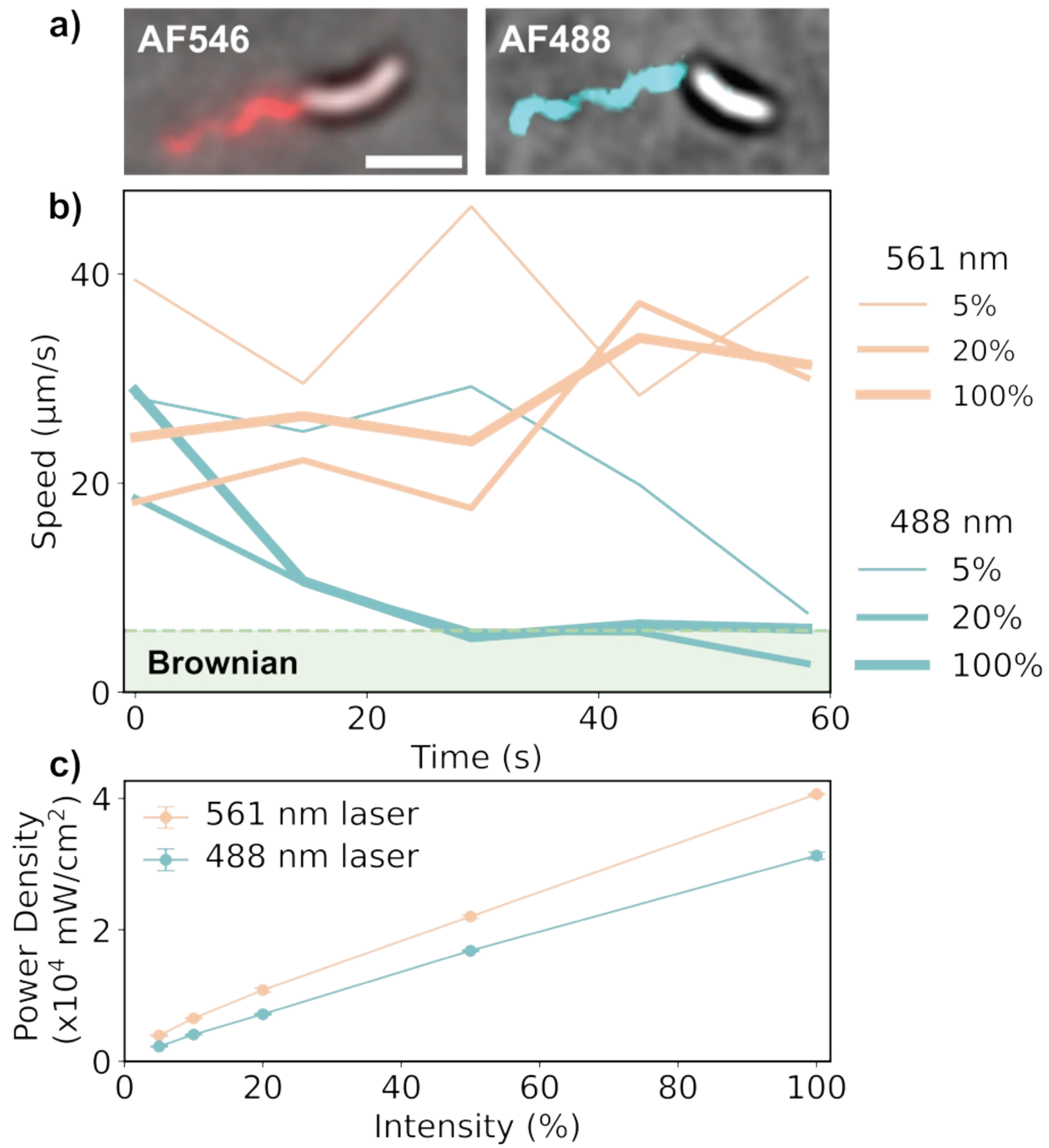


**Figure S9**: **Imaging and laser intensity analysis. a)** Merged images of brighfield swimmer and its labeled flagella on an agarose pad. Left: Flagella labeled with AF546 dye. Right: Flagella labeled with AF488 dye. Scale bar: 2 µm. **b)** Wild-Type swimmer without labeling, trapped inside a µ-Trap. Speed vs. Time under different laser intensities and wavelengths. **c)** Measured laser power density and corresponding intensity values. (561 nm: 5% - $3.9 \times 10^3$ mW/cm$^2$, 20% - $1.1 \times 10^4$ mW/cm$^2$, 100% - $4.1 \times 10^4$ mW/cm$^2$) (488 nm: 5% - $2.3 \times 10^3$ mW/cm$^2$, 20% - $7.2 \times 10^3$ mW/cm$^2$, 100% - $3.1 \times 10^4$ mW/cm$^2$).

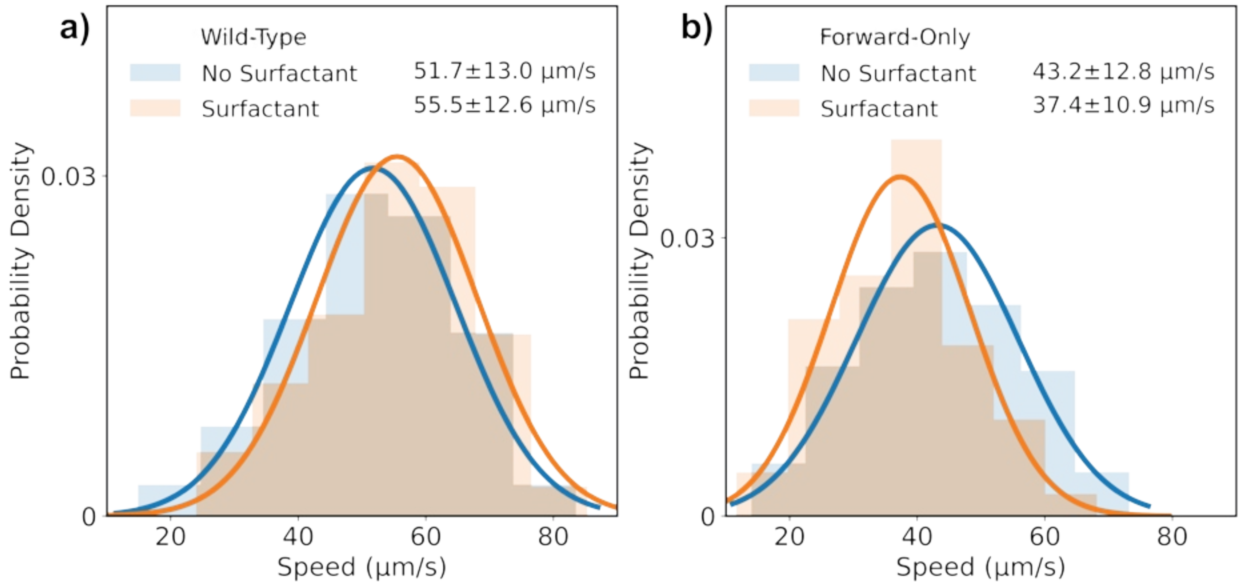


**Figure S10**: **No significant effect of experimental surfactant concentration (0.002 wt% Pluronic® F108) on swimmer speed. a)** Wild-Type strain: N = 258 (no surfactant), N = 263 (with surfactant). **b)** Forward-Only strain: N = 249 (no surfactant), N = 262 (with surfactant).

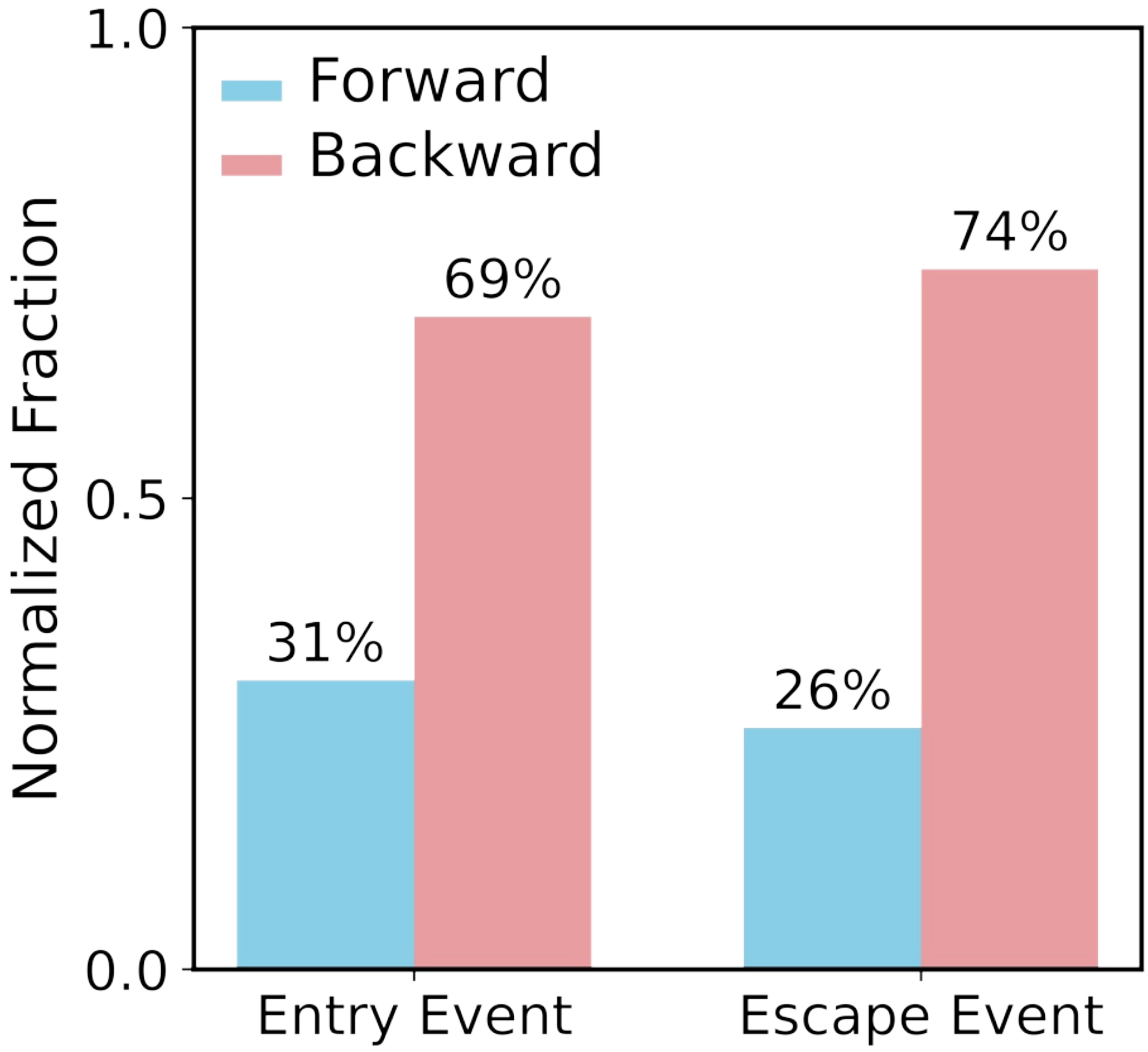


**Figure S11**: **Fraction of entry and escape events normalized by the forward/backward motion bias.** A total of 60 forward entry and 70 backward entry events were recorded. The fraction of backward-moving bacteria outside the µ-Trap was approximately 34%. For escape events, 57 forward escapes (combining Forward I and Forward II) and 89 backward escapes were observed. The fraction of backward-moving bacteria inside the µ-Trap was approximately 35%.

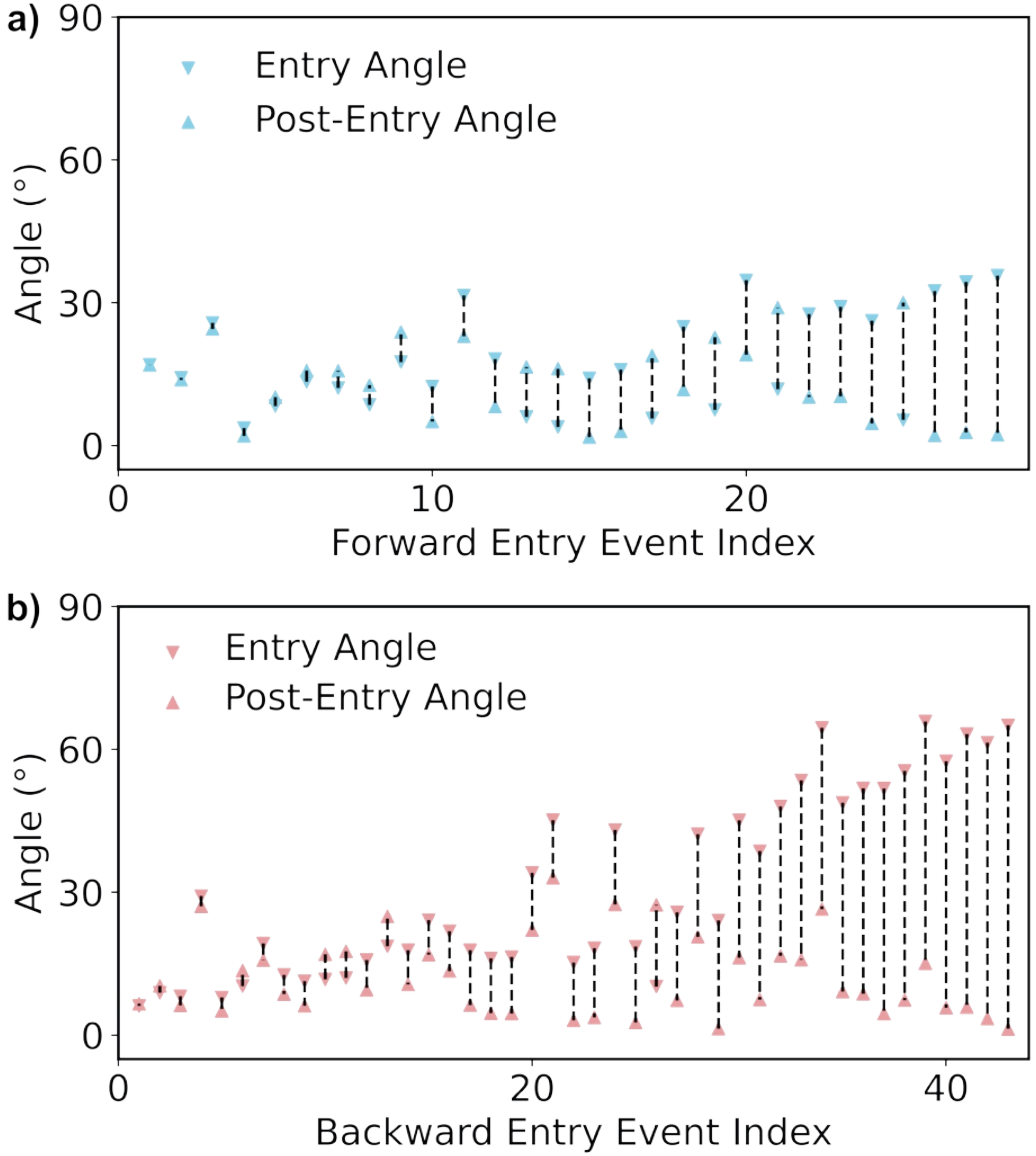


**Figure S12**: **Changes in entry angle and post-entry angle. a)** Forward entry (N = 28). **b)** Backward entry (N = 43).

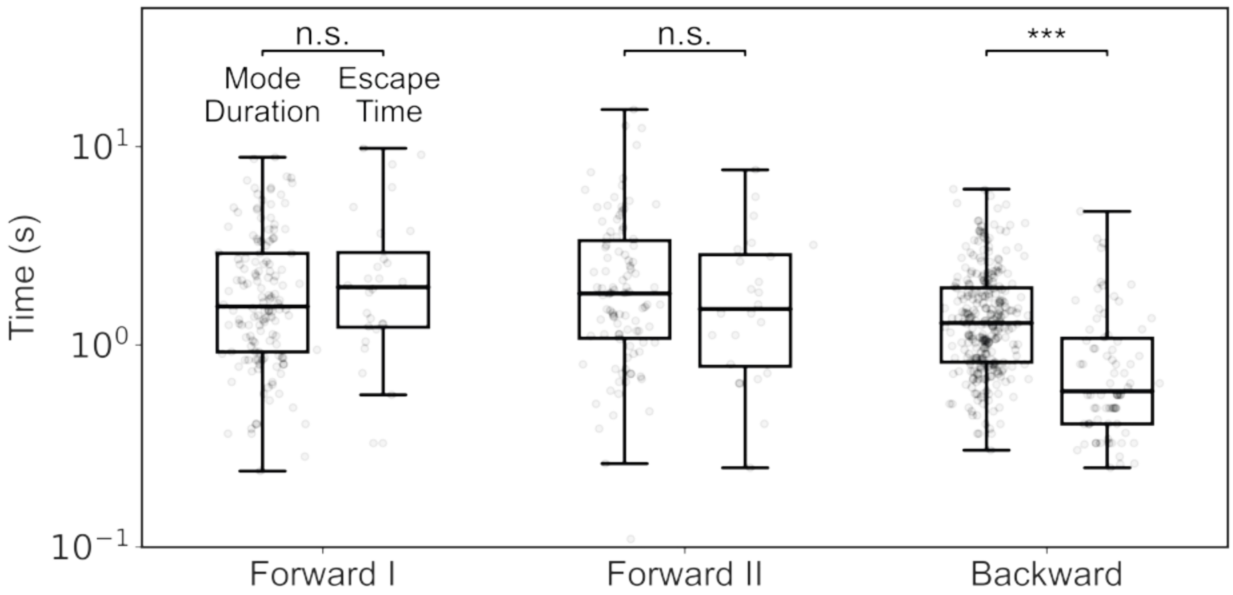


**Figure S13**: **Comparison between mode duration and escape time after the last switch.** This comparison shows that although the Backward mode duration is shorter, the Backward escape time is not limited by the Backward mode duration. Mode durations exclude the start and end time durations. (Forward I: mode duration N = 140, escape time N = 28, $p = 0.4398$, not significant; Forward II: mode duration N = 98, escape time N = 24, $p = 0.4320$, not significant; Backward: mode duration N = 268, escape time N = 74, $p < 0.001$, ***)

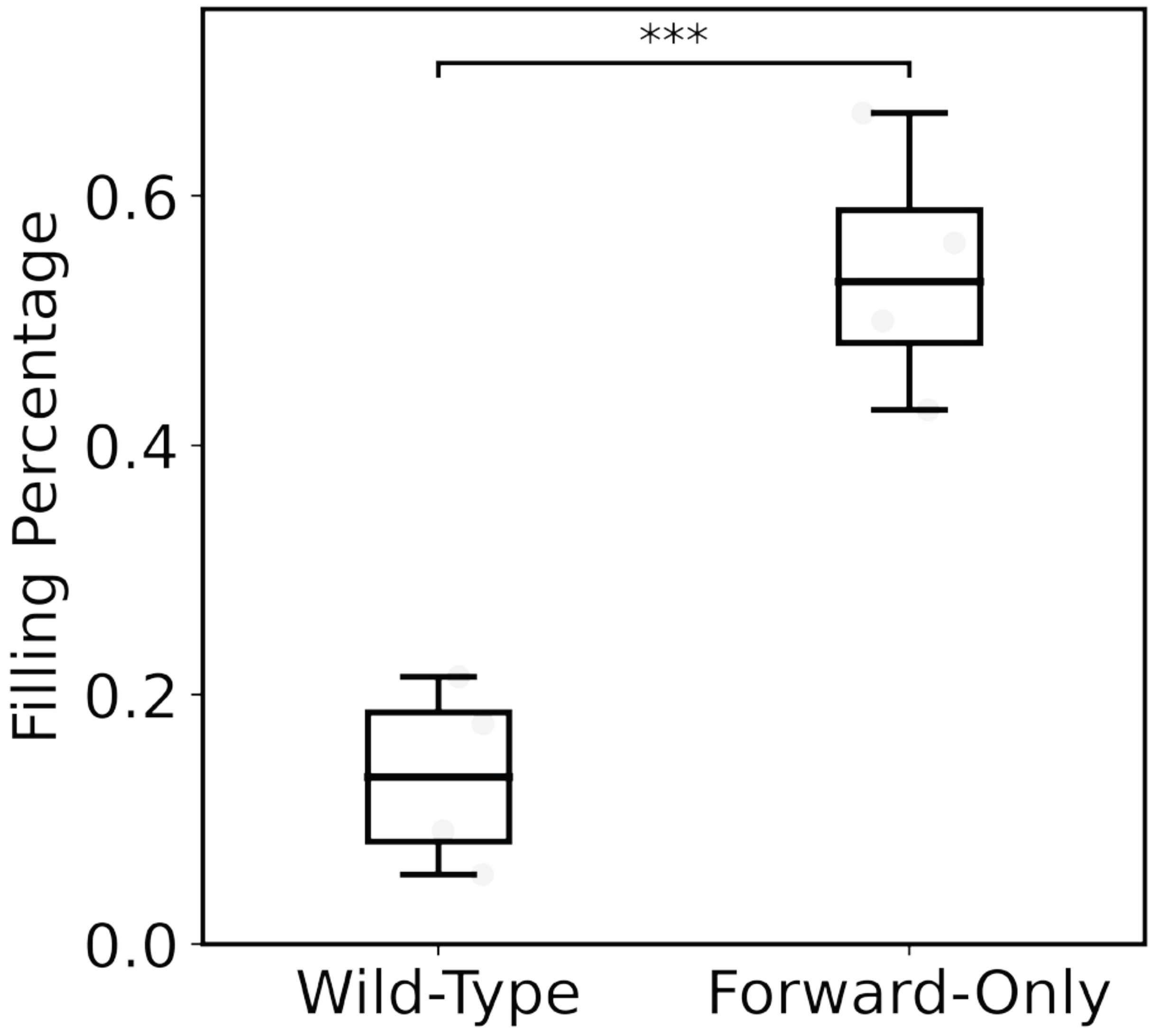


**Figure S14**: **Filling percentage after 1 hour post addition.** Four fields of view were compared (N = 4), each containing 11-21 µ-Traps. (***: $p$ < 0.001)

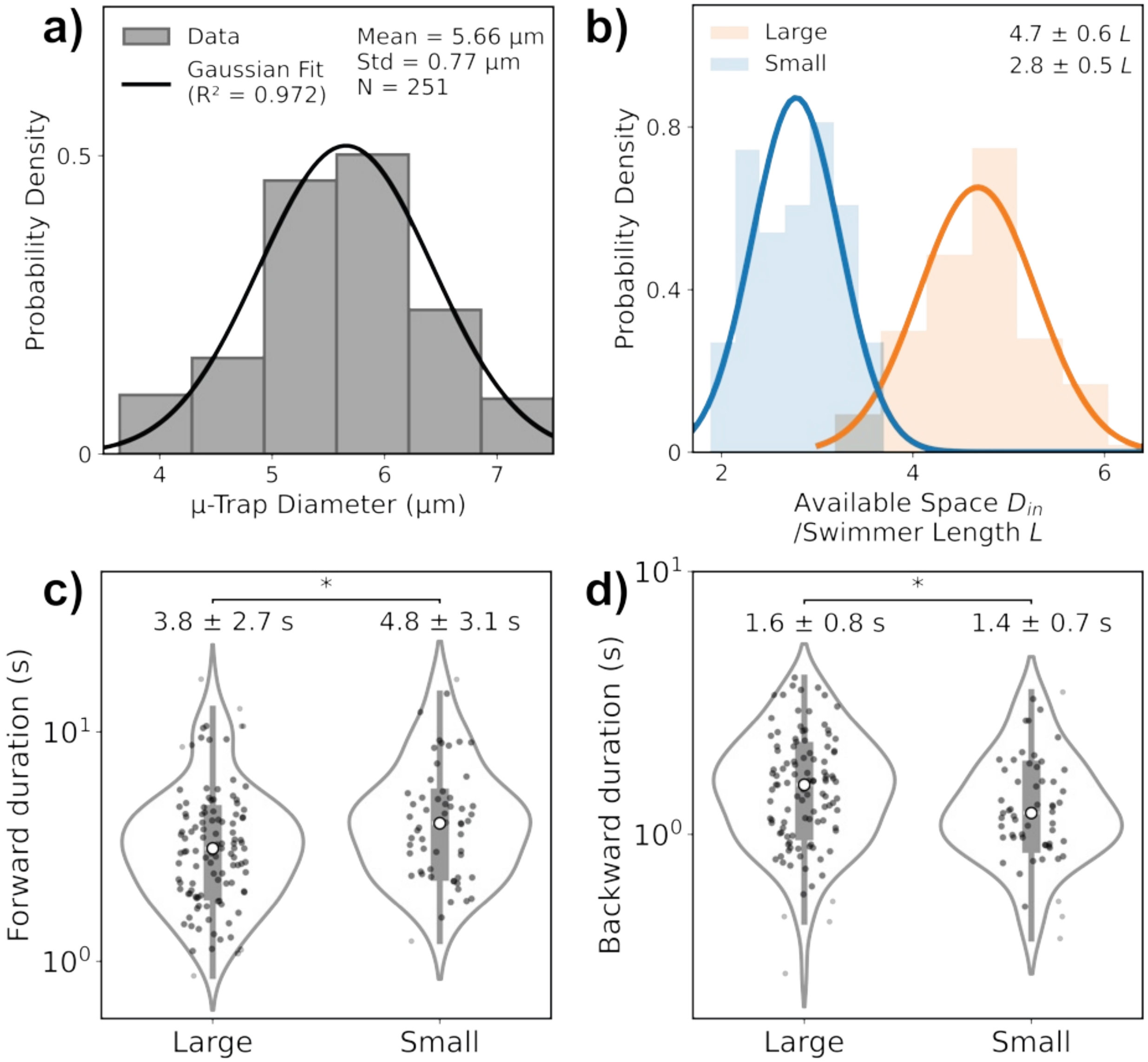


**Figure S15**: **Quantification of geometry and motion durations in large and small µ-Traps. a)** Distribution of diameters for small µ-Traps (N = 251). **b)** Distribution of relative available space in large and small µ-Traps. Relative space is defined as the ratio between the inner diameter of the µ-Trap ($D_{in}$) and the swimmer body length ($L$). For panels (**b–d**): N = 108 for large µ-Traps, N = 57 for small µ-Traps. **c)** Violin plot comparing forward swimming durations in large and small µ-Traps, showing significantly longer forward swimming durations in small µ-Traps (*, $p$ = 0.0131). **d)** Violin plot comparing backward swimming durations in large and small µ-Traps, showing significantly shorter backward swimming durations in small µ-Traps (*, $p$ = 0.0312). In panels (**c-d**), white circles indicate median values; mean ± standard deviation are shown above.

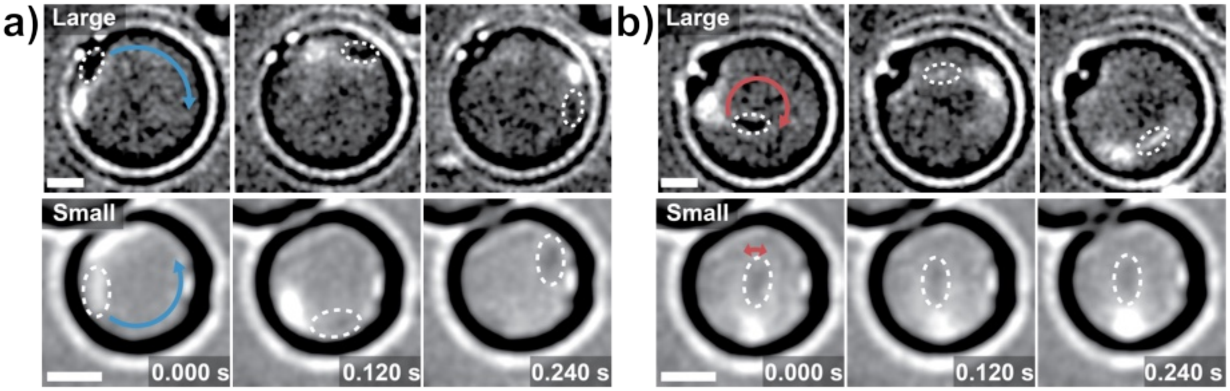


**Figure S16**: **Flagella-labeled time-lapse images illustrating the effect of confinement on bacterial swimming behavior.** Blue arrows indicate forward motion; red arrows indicate backward motion; white dashed circles outline cell bodies. Scale bars: 2 µm. **a)** Unimpeded forward swimming in both large (top) and small (bottom) µ-Traps. **b)** Suppressed backward motion in small µ-Traps (bottom), unimpeded backward motion in large µ-Traps (top).

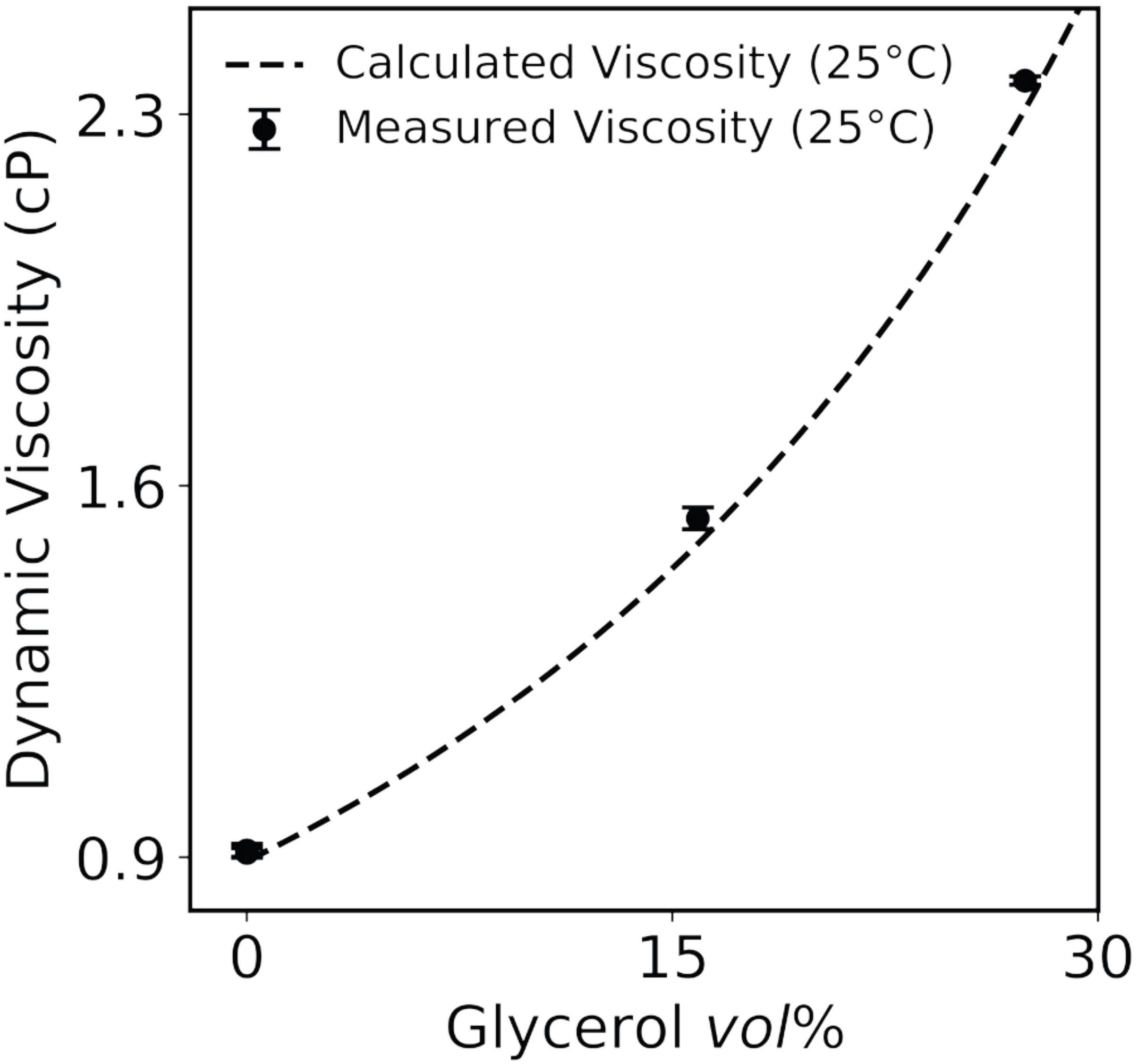


**Figure S17**: **Dynamic viscosity of glycerol-M2 mixture vs. glycerol volume fraction and the calculated viscosity of glycerol-water mixture.** Data shown were measured from three repetitions. Temperature was controlled at 25°C.

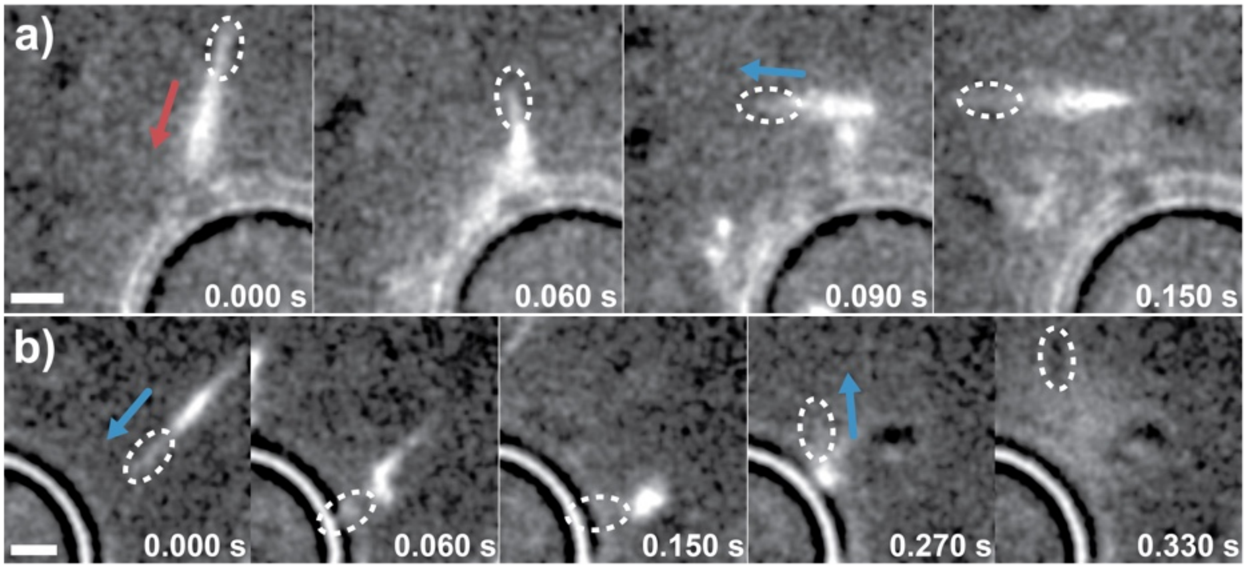


**Figure S18**: **Flagellum-specific mechanical gating of backward motion. a**) Time-lapse images of a flagella-labeled Caulobacter cell swimming backward toward a µ-Trap. Upon flagellar contact with the surface, the cell immediately switches direction. **b)** Time-lapse images of a flagella-labeled cell swimming forward into a surface. Despite collision with the cell body, no directional switch occurs. White dashed circles indicate the cell body. Scale bars: 2 µm. Red arrows: backward motion. Blue arrows: forward motion.

**Table S1**: **Resources and Materials**

| **1. *Caulobacter crescentus* Strains** | **Description** | **Source or Reference** |
| --- | --- | --- |
| SS065 | NA1000 | Evinger et al. (*38*) |
| SS494 | NA1000 *fljK*$^{T103C}$ | This study |
| SS598 | NA1000 Δ*cheR1* Δ*cheR2* Δ*cheR3* | MT293, Briegel et al. (*39*) |
| **2. Plasmids** | | |
| SS490 | pNPTS139 *fljK*$^{T103C}$ | YB6566, Berne et al. (*36*) |
| **3. Chemical and Other Materials** | | |
| 3-(Trimethoxysilyl)propyl methacrylate (TPM) | M6514-25ML | Sigma-Aldrich |
| Polystyrene beads | PS_1.3µm_16.3wt% | Sacanna Lab |
| Tetrahydrofuran | T425-4 | Fisher Scientific |
| Pluronic® F108 | 07059 | Sigma-Aldrich |
| Alexa Fluor™ 488 C5 Maleimide | A10254 | Invitrogen |
| Alexa Fluor™ 546 C5 Maleimide | A10258 | Invitrogen |
| Nile Blue A | N0766 | Sigma-Aldrich |
| DMSO | D12345 | Invitrogen |
| Percoll® | P4937-500mL | Sigma-Aldrich |
| Hydrogen peroxide | H325-500 | Fisher Scientific |
| Agarose | 16520050 | Invitrogen |

Continued on next page

Table S1 – continued from previous page

| 3. Chemical and Other Materials | Description | Source or Reference |
|---|---|---|
| 384 well glass bottom plate | P384-1.5H-N | Cellvis |
| Microscope slide | 12-550-A3 | Fisher Scientific |
| Microscope coverslip | 12-541-029 | Fisher Scientific |
| Grade A-E glass microfiber filters | 1229B35 | Thomas Scientific |
| Petri dish | D210-16 | Thomas Scientific |
| **4. Software and Algorithms** | | |
| FIJI | RRID:SCR_002285 | Schindelin et al. (*40*) |
| TrackMate | ImageJ plugin | Ershov et al. (*41*) |
| ilastik | RRID:SCR_015246 | Berg et al. (*42*) |
| MicrobeJ | RRID:SCR_023914 | Ducret et al. (*43*) |
| Adobe Illustrator | RRID:SCR_010279 | Adobe software |
| Adobe Premiere Pro | RRID:SCR_021315 | Adobe software |

**Caption for Movie S1. 3D tracking of bacteria in µ-Traps.** Animated version of Fig. 2c-d.

**Caption for Movie S2. Bacterial motion modes in µ-Traps.** Part 1: Brightfield imaging. Part 2: Flagella labeling.

**Caption for Movie S3. Rapid reduction in motility under 488 nm laser excitation.**

**Caption for Movie S4. Bacterial entry into µ-Traps.** Part 1: Brightfield imaging. Animated version of Fig. 3a. Part 2: Flagella labeling.

**Caption for Movie S5. Bacterial escape from µ-Traps.** Part 1: Brightfield imaging. Animated version of Fig. 3c. Part 2: Flagella labeling.

**Caption for Movie S6. Trapping efficiency after 1 hour post-addition.** Wild-Type vs. Forward-Only.

**Caption for Movie S7. Increased backward-swimming bacteria near the surface.**

**Caption for Movie S8. Impeded backward motion in small µ-Traps.**

**Caption for Movie S9. Nearly no backward-swimming bacteria at high viscosity.**

**Caption for Movie S10. Flagellar contact triggers an instant directional switch; body contact does not.**
